# Cell-type-specific splicing of transcription regulators and *Ptbp1* by *Rbfox1/2/3* in the developing neocortex

**DOI:** 10.1101/2024.09.09.612108

**Authors:** Xiangbin Ruan, Kaining Hu, Yalan Yang, Runwei Yang, Elizabeth Tseng, Bowei Kang, Aileen Kauffman, Rong Zhong, Xiaochang Zhang

**Affiliations:** Department of Human Genetics, The University of Chicago, Chicago, IL 60637, USA; Pacific Biosciences, Menlo Park, CA 94025, USA; The Neuroscience Institute, The University of Chicago, Chicago, IL 60637, USA

**Keywords:** long-read RNA-Seq, full-length isoforms, NeuN, unproductive splicing, multiplexed CRISPR editing, neuronal migration

## Abstract

How master splicing regulators crosstalk with each other and to what extent transcription regulators are differentially spliced remain unclear in the developing brain. Here, cell-type-specific RNA-Seq of the developing neocortex uncover that transcription regulators are enriched for differential splicing, altering protein isoforms or inducing nonsense-mediated mRNA decay. Transient expression of Rbfox proteins in radial glia progenitors induces neuronal splicing events preferentially in transcription regulators such as *Meis2* and *Tead1*. Surprisingly, Rbfox proteins promote the inclusion of a mammal-specific alternative exon and a previously undescribed poison exon in *Ptbp1*. Simultaneous ablation of *Rbfox1/2/3* in the neocortex downregulates neuronal isoforms and disrupts radial neuronal migration. Furthermore, the progenitor isoform of *Meis2* promotes *Tgfb3* transcription, while the *Meis2* neuron isoform promotes neuronal differentiation. These observations indicate that transcription regulators are differentially spliced between cell types in the developing neocortex.

## Introduction

Alternative pre-mRNA splicing is enriched in the brain and occurs between developmental stages, brain regions, and cell types(*1–7*). Alternative splicing is modulated by cis-regulatory sequences and their associated RNA-binding proteins(*8–12*). Human mutations have been reported to shift splice isoforms and cause neurodevelopmental disorders(*13–16*). Dysregulated splicing networks have also been reported as one of the converging mechanisms for autism pathogenesis(*17–19*). These observations underscore the importance of alternative splicing in brain development.

The neocortex is formed by millions of neurons in mice and billions in humans(*20, 21*), and the generation of cortical neurons requires molecular balances between progenitor self-renewal and differentiation(*22–24*). In mice, radial glial cells (RGCs) start to produce neurons and intermediate progenitor cells (IPCs) at embryonic day 11.5 (E11.5), and the IPCs further divide to form cortical projection neurons between E11.5 and E18.5(*25, 26*). Differential expression of transcription regulators plays an important role in cortical development(*27*). Interestingly, alternative splicing enables the neuronal transcription repressor *Rest* to express its functional form in neural progenitors but its non-functional isoform in differentiated cells(*28*). Recent comparisons of isolated cortical RGCs and non-RGCs uncovered hundreds of alternative exons and their roles in shaping neurogenesis by remodeling the cytoskeleton(*13, 29*). However, to what extent transcription regulators are differentially spliced between cell types remains largely unexplored in the developing neocortex.

Three Rbfox proteins, including the pan-neuronal marker Rbfox3/NeuN, have been reported as a hub for autism pathogenesis(*17, 18*), but it has been challenging to uncover their functions in the neocortex due to genetic redundancy. While *Rbfox1* and *Rbfox3* knockout mice are susceptible to seizures, single *Rbfox1*, *Rbfox2*, or *Rbfox3* knockout mice display relatively normal cortical morphology and limited splicing changes when compared to CLIP-Seq findings(*30–33*). Prominently, the Rbfox1 protein level is increased in *Rbfox2* knockout mouse brains(*30*), while the Rbfox2 level is increased in *Rbfox1* knockout(*31*). Thus, the molecular and physiological functions of Rbfox proteins in cortical development await further investigation.

*Ptbp1* is a master splicing regulator, and *Ptbp1* knockout led to lethal hydrocephalus in mice(*13, 34*). *Ptbp1* suppresses neuronal differentiation in nonneural cells and neural progenitors(*35–37*), but how it is confined in proliferating cells remains unclear. Ptbp1 has been reported to express in RGCs and suppress neuronal exon inclusion, while Rbfox proteins in neurons promote neuronal exon insertion (*3, 13, 38*). It remained unknown whether *Rbfox* and *Ptbp* genes directly interact in the brain. Furthermore, the mammal-specific alternative exon8 in *Ptbp1* (exon9 of human *PTBP1*) regulates an evolutionarily unique set of splicing events(*39, 40*). However, it is unclear how the *Ptbp1* mammal-specific splicing is regulated.

In this study, we compared cell populations using bulk and single-cell approaches and showed that transcription regulators were enriched for alternative splicing during cortical neurogenesis. Transient Rbfox expression in RGCs promoted neuronal splice forms for dozens of transcription regulators, while simultaneous ablation of *Rbfox1/2/3* in the brain disrupted radial neuronal migration and decreased neuronal splice isoforms. Rbfox proteins directly regulate *Meis2* isoforms that show different functions in cortical development. Rbfox expression also promoted the inclusion of the *Ptbp1* mammal-specific exon8 and a previously unannotated poison exon that downregulated *Ptbp1* expression. Thus, Rbfox proteins antagonize *Ptbp1* through unproductive splicing and coordinate isoform switching of transcription regulators in cortical development.

## Results

### Transcription regulators are differentially spliced in the developing neocortex

We sorted EGFP-positive and -negative cells from the E14.5 mouse dorsal forebrains using the *Tg(Tubb3:EGFP)* and *Tg(Eomes:EGFP)*(*41*) BAC transgenic lines which express EGFP in neurons or a mixture(*13*) of IPCs and immature neurons, respectively (Fig. 1a, Fig. S1a). We performed RNA-Seq for sorted cell population and identified differentially expressed genes: the RGC marker *Nes* was highly enriched in *Eomes:EGFP*(-) cells, the IPC markers *Eomes* and *Btg2* were enriched in *Eomes:EGFP*(+) samples, and the neuronal markers *Dcx* and *Stmn2* were enriched in the *Tubb3:EGFP*(+) cells (Fig. 1b). Interestingly, we found that *Eomes:EGFP*(+) cells expressed a unique set of genes such as *Mfap4*, *Egr1*, *Sstr2* and *Cdh8* (Fig. 1b). These results indicate that the isolated *Eomes:EGFP*(-), *Eomes:EGFP*(+) and *Tubb3:EGFP*(+) cells are enriched for RGCs, IPCs, and cortical neurons, respectively.

**Figure 1.**
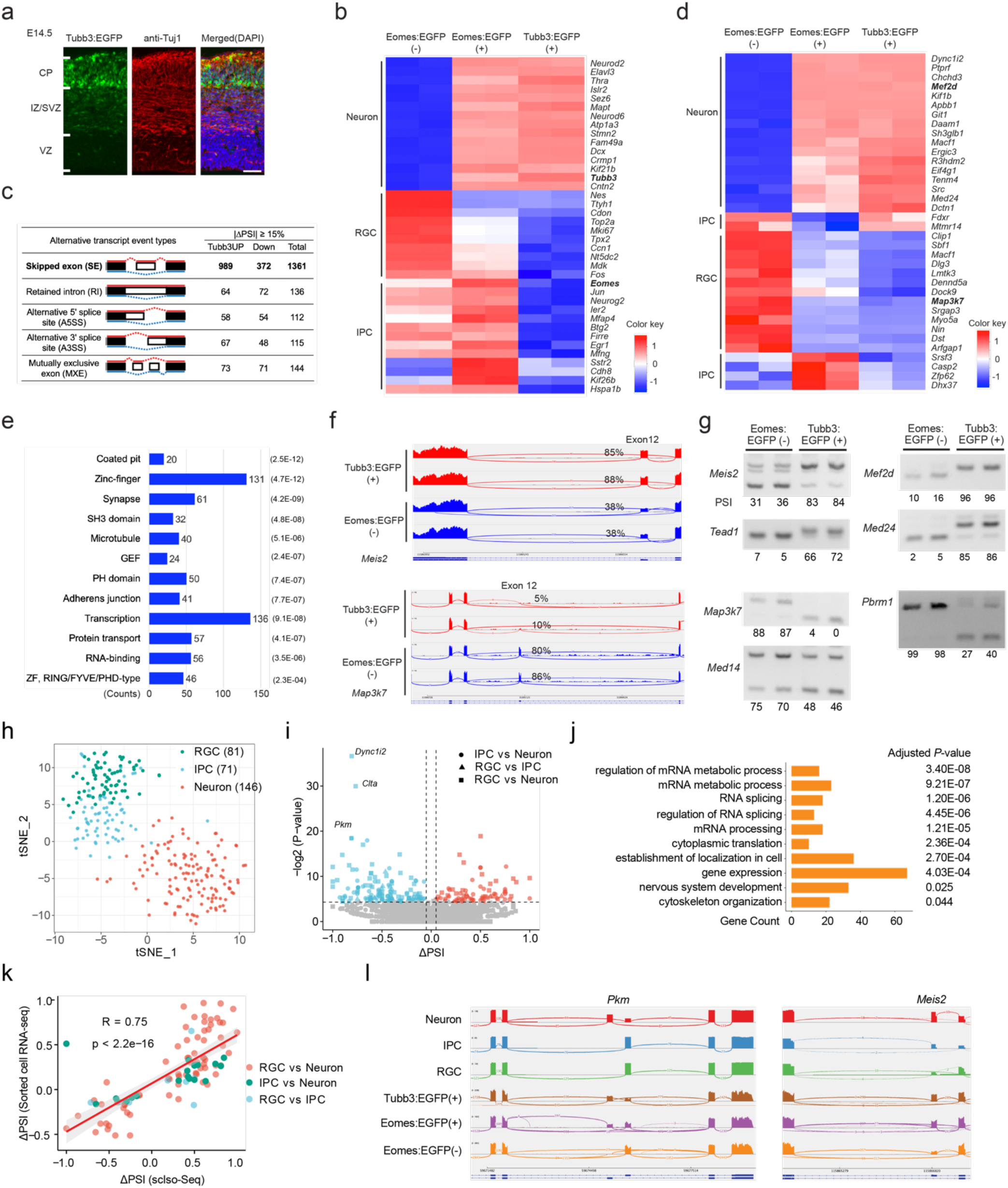
Transcription regulators are differentially spliced in the developing neocortex. **a.** *Tubb3:EGFP* labels cortical neurons in E14.5 mouse dorsal forebrain, scale bar, 100μm. **b.** Heatmap of mRNA levels in *Eomes:EGFP*(*-*), *Eomes:EGFP(+)*, and *Tubb3:EGFP(+)* cells showing RGC-, IPC-, and neuron-specific genes, respectively. **c.** RNA-Seq of *Tubb3:EGFP*(+) cells and *Eomes:EGFP*(-) cells uncovered alternative splicing changes, PSI = percentage spliced in (|ΔPSI| >= 15%, FDR < 0.05); **d.** Heatmap of PSI values showing alternative splicing events specific to indicated cell populations. **e.** Transcription regulators and synaptic transmission-related genes are highly enriched for differentially spliced genes. Top-to-bottom ranked by enrichment scores. The numbers of associated gene counts and FDR (numbers in parentheses) are indicated. **f.** Sashimi plot showing neuron-enriched (*Tubb3:EGFP*(+)) exon12 in transcription regulator *Meis2*. **g.** RT-PCR validation of neuron-enriched (*Tubb3:EGFP*(+)) exons in *Meis2*, *Tead1, Mef2d* and *Med24*, and RGC-enriched (*Eomes:EGFP*(-)) exons in *Map3k7, Med14* and *Pbrm1*. numbers showing PSI values. All RT-PCR splicing analyses were statistically significant by t-test (p < 0.05, the details of statistical comparisons for each experiment are included in Table S2). **h.** A tSNE plot based on full-length, single-cell transcripts where each point represents a cell, color coded by cell types. **i.** Volcano plot showing the exons that are differentially spliced between RGCs, IPCs, and neurons. **j.** Gene ontology analysis of genes with differentially spliced exons (DAVID 6.8). **k.** A scatter plot showing the ΔPSI values of differentially spliced exons are highly correlated between scIso-seq and bulk RNA-seq results. **l.** Sashimi plots showing differentially spliced exons of *Pkm* and *Meis2* in scIso-Seq (Neuron, IPC, RGC) and sorted bulk samples (Figure 1). See also Figure S1.

We uncovered cell-type-specific alternative splicing events between *Eomes:EGFP*(-), *Eomes:EGFP*(+), and *Tubb3:EGFP*(+) cells (the absolute difference of percentage spliced in values or |ΔPSI| > 0.15, FDR < 0.05, details in Methods, Fig. 1c). Cassette/skipped exons (SEs) formed the largest group (1361 exons) and we focused on them hereafter. We found that the poison exon in master splicing regulator *Srsf3*(*42*), as well as the reported *Srsf3* target exon in the Caspase2 gene *Casp2*(*43*), showed higher inclusion in *Eomes:EGFP*(+) cells than in *Tubb3:EGFP*(+) neurons (Fig. 1d), suggesting IPC-specific splicing events.

We performed gene ontology analysis and found coated pit, zinc-finger and synapse were the most significantly enriched terms (Fig. 1e). Interestingly, zinc-finger (131) and transcription regulators (136) had the largest number of genes that were differentially spliced during neurogenesis (Fig. 1e). For example, *Mef2d* exon8 and *Meis2* exon12 showed higher inclusion in *Tubb3:EGFP*(+) neurons, while *Map3k7* exon12 was enriched in *Eomes:EGFP*(-) progenitor cells (Fig. 1f-1g, Fig. S1c). We selected 22 most differentially spliced exons in transcription regulators and validated all of them with RT-PCR except for *Arntl* where the primers failed to detect the expected product (Fig. 1g and Fig. S1b-S1c).

We integrated droplet-based single-cell RNA-Seq (scRNA-Seq)(*44*) with long-read sequencing (scIso-Seq) to ascertain cell-type-specific splicing changes in the E14.5 mouse dorsal forebrain. After single-cell capture and SMART-PCR, the full-length cDNA library was split into two fractions: one fraction was tagmented and further processed for short-read sequencing and cell type identification (Fig. 1h and S1d), and the untagmented cDNA fraction was subjected to PacBio Sequel2. We obtained 1.8 million long reads after the identification of UMIs, cell barcodes, and the removal of PolyA tail and concatemers. We assigned 0.64 million long reads to annotated cell types (Fig. S1e). We grouped long reads from each cell type as pseudo bulk long-read samples and identified differentially spliced exons between RGCs, IPCs, and neurons using FLAIR(*45*) (Fig. 1i). Gene ontology analysis indicated that “mRNA processing” was highly enriched, and the “gene expression” term showed the highest number of genes (Fig. 1j).

Importantly, the differential exons uncovered from the long-read comparisons are concordant with those identified from FACS-Sorted samples enriched for RGCs, IPCs, and neurons (Fig. 1k, R=0.75, P < 2.2e-16). For example, the alternative exons between RGCs and neurons in *Meis2, Map3k7*, and *Pkm* were validated by scIso-Seq (Fig. 1l); the ultraconserved *Srsf3* exon significantly enriched in *Eomes:EGFP(+)* cells was also significantly included in scIso-Seq IPCs (Fig. S1f). In summary, the single-cell long-read results uncover splicing changes between RGCs, IPCs, and neurons during cortical neurogenesis and cross-validate findings from the FACS-sorted cell populations.

### Alternative exons in transcription regulators are predicted to affect protein expression

Among alternative exons in transcription regulators, 16 SEs were predicted to cause nonsense-mediated mRNA decay (AS-NMD)(*46*) when inserted (NMD_in, 3 events) or excluded (NMD_ex, 13 events, Fig. 2a). In addition, 17 SEs were in 5’ untranslated regions (UTRs), 7 SEs contained start codons, 8 SEs altered open reading frames (ORFs), and 54 SEs preserved ORFs. We further analyzed ORF-preserving SEs and found that 17 SEs were inside protein domains, with 6 additional SEs switching protein domains on or off (Fig. 2b-2d, Fig. S2a-S2b). For example, the inclusion of neuronal exon12 in *Meis2* led to the truncation of its C-terminal end (Fig. S2c). Interestingly, the inclusion of *Crtc2* exon 13N was enriched in neurons and increased during cortical development, leading to a premature stop codon in the TORC-C domain that was predicted to cause NMD (Fig. 2e-2g). To validate predicted NMD exons, we treated primary hippocampal neurons (1 day cultured *in vitro*, DIV1) with cycloheximide (100 μg/ml, 5 hours) to suppress protein translation and NMD. Among predicted NMD exons, NMD_in exons in *Crtc2* and *Whsc1* were significantly increased after cycloheximide treatment (Fig. 2h-2i), while three NMD_ex exons (*Ccar2*, *Taf1a*, *Ylpm1*) were significantly decreased (Fig. 2j). These results indicate that alternative splicing of transcription regulators alters protein isoforms or gene expression levels through NMD.

**Figure 2.**
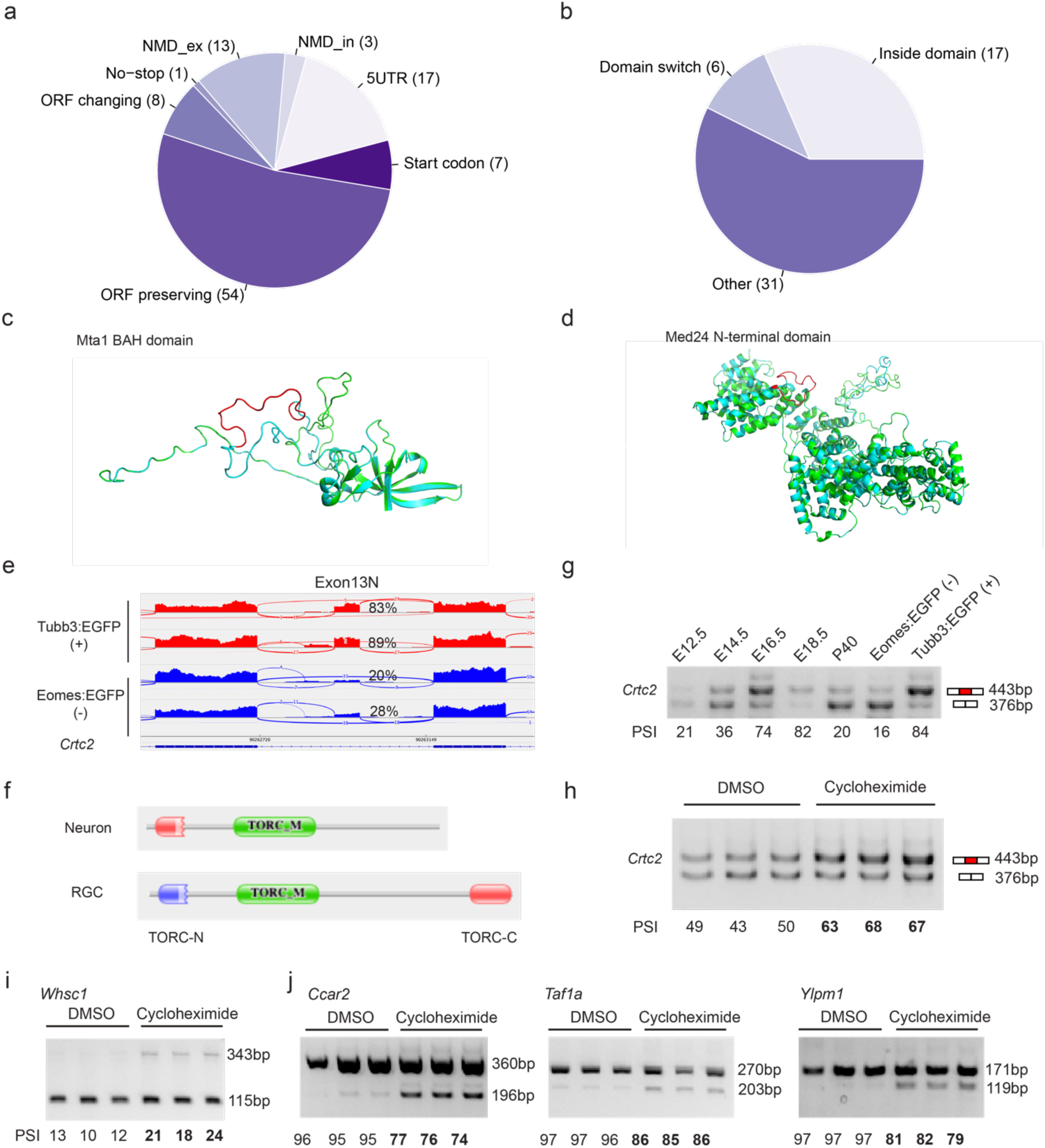
Alternative exons are predicted to regulate protein expression. **a.** Alternative exons in transcription regulators were predicted to cause nonsense-mediated mRNA decay when included (NMD_in) or excluded (NMD_ex), or alter protein sequences. ORF changing refers to SEs that were predicted to change the open reading frame while not inducing NMD. ORF preserving refers to SEs that were predicted to be in-frame. **b.** 54 ORF-preserving alternative exons (Fig. 2A) in transcription regulators were predicated for their effects on modular protein domains. “Inside domain” refers to SEs that are part of a predicted protein domain; “Domain switch” means that one of the isoforms has a predicted protein domain, but the other splice isoform of the same gene does not. **c.** 3-D protein structure (SWISS-MODEL) showing that a 51-bp alternative exon in *Mta1* alters the BAH domain. The alternative exon is in red. **d.** Alternative splicing of a 57-bp exon (red) in *Med24* was predicted to affect the N-terminal domain. **e.** Sashimi plot showing alternative splicing of *Crtc2* that was predicted to introduce premature stop codons in *Tubb3:EGFP*(+) neurons. **f.** The inclusion of the *Crtc2* neuronal exon was predicted to introduce premature stop codons in front of the TORC-C domain. **g.** Inclusion of the *Crtc2* exon13N was increased during embryonic neocortex development. **h.** RT-PCR results showing that cycloheximide (CHX) treatment of DIV1 primary hippocampal neurons increased *Crtc2* exon13N inclusion. **i.** Cycloheximide (CHX) treatment led to increased inclusion of the *Whsc1* exon. **j.** CHX treatment decreased inclusion of exons in *Ccar2, Taf1a*, and *Ylpm1*. See also Figure S2.

### *Rbfox1/2/3* display overlapping and variable expression patterns in the developing neocortex

Canonical Ptbp and Rbfox binding motifs were highly enriched in the upstream or downstream introns of neuron-specific exons (Fig. S3a). *Rbfox1/2/3* have been reported to express in cortical neurons, but whether they share the same expression pattern among cortical cell types remained unclear. Surprisingly, *Rbfox3* showed enriched expression in FACS-sorted *Eomes:EGFP*(+) cells, suggesting that *Rbfox3* is transcribed in immature neurons and possibly in Eomes-positive cells (Fig. 3a). We analyzed scRNA-Seq results from E14.5 cortical cells(*44*) and plotted *Ptbp1* and *Rbfox1/2/3* expression (Fig. 3b-3c). As expected, *Ptbp1* showed higher expression in RGCs and diminished in IPCs. In contrast, *Rbfox3* transcripts showed higher expression in immature neurons, and *Rbfox2/1* mRNA showed their peak expression in mature neurons (Fig. 3b-3c). Further analysis of an independent E14.5 scRNA-Seq dataset(*47*) confirmed the mRNA expression patterns of *Rbfox1/2/3* (Fig. S3b).

**Figure 3.**
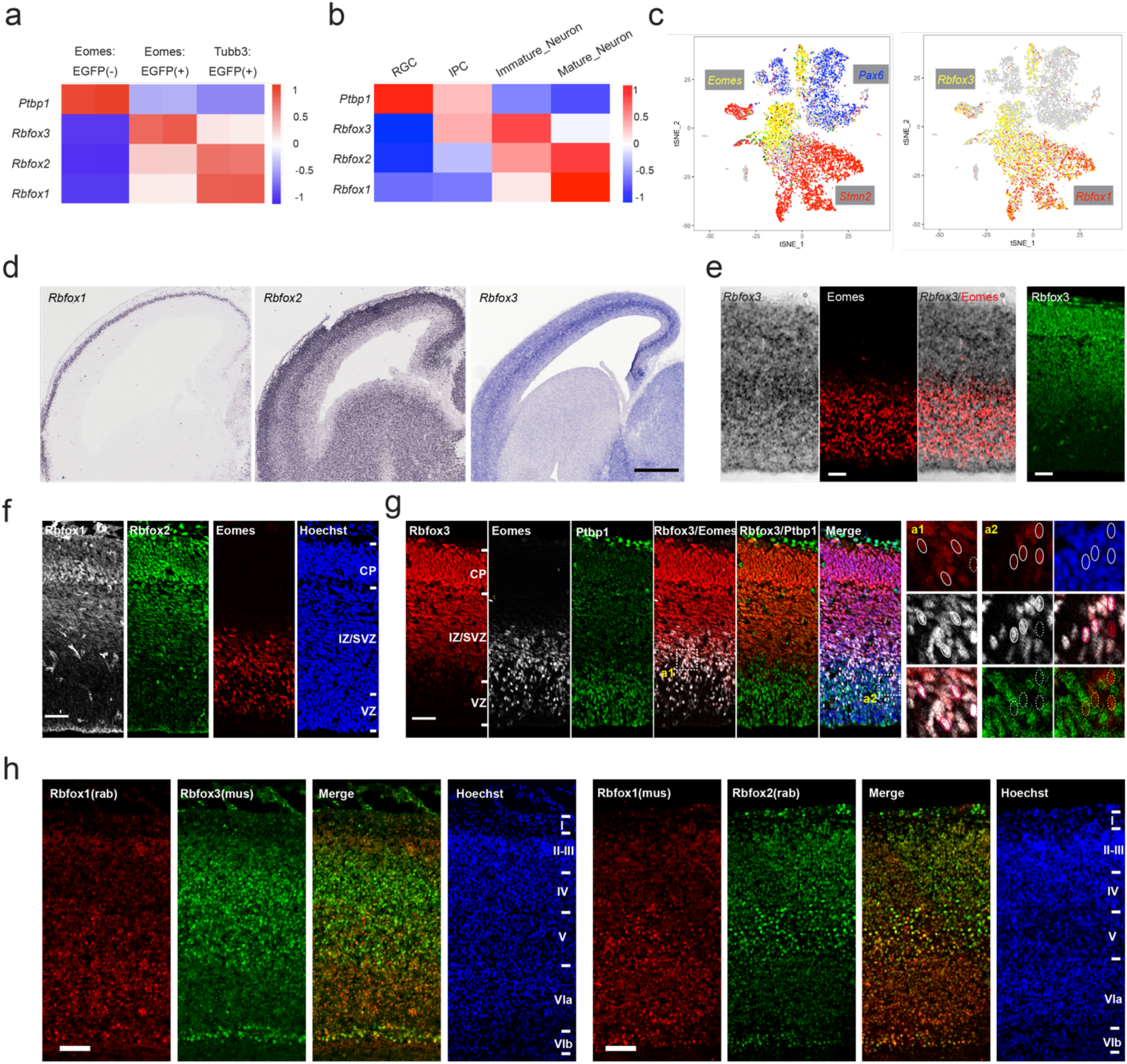
Differential *Rbfox3/2/1* expression in the developing neocortex. **a.** *Rbfox3/2/1* and *Ptbp1* mRNA levels in bulk RNA-Seq data of E14.5 *Eomes:EGFP*(-), *Eomes:EGFP*(+) and *Tubb3:EGFP*(+) cells. Red-blue represents high-low gene expression. **b.** Transcription levels of *Ptbp1* and *Rbfox3/2/1* in E14.5 RGCs, IPCs, immature and mature neurons. Re-analysis of our published scRNA-Seq data(*44*). **c.** Feature plots of E10.5-E18.5 scRNA-Seq(*44*) data showing *Rbfox3* expression in immature neurons and a fraction of *Eomes*-positive cells. **d.** RNA ISH results of E14.5 sagittal brain sections showing the expression of *Rbfox3* (this study, with a probe against the coding sequence) in comparison to *Rbfox1* and *Rbfox2* (adapted from Genepaint(*48*)). Scale bar, 400μm. **e.** Eomes/Tbr2 immunostaining and *Rbfox3* RNA ISH co-labeling showed overlapping signals in the SVZ/IZ. Rbfox3 immunostaining was performed on an adjacent section to the RNA ISH one. Scale bar, 50μm. **f.** Co-Immunostaining of Rbfox1 and Rbfox2 in the E14.5 dorsal forebrain. CP: cortical plate; VZ: ventricular zone. Scale bar, 100μm. **g.** Co-immunostaining results of Rbfox3 with Eomes and Ptbp1 in the E14.5 mouse dorsal forebrains. Rbfox3 was expressed in 39% of Eomes-positive cells while overlapped with less than 2.5% of Ptbp1-positive cells. Scale bar, 100μm. **h.** Co-immunostaining results of Rbfox1 (Sigma-Aldrich HPA040809, rabbit) and Rbfox3 (Millipore MAB377, mouse), or Rbfox1 (Millipore MABE159, mouse) and Rbfox2 (Bethyl A300-864A, rabbit) in the dorsal forebrains of P0 mice. Scale bar, 100μm. See also Figure S3.

We performed RNA in situ hybridization (ISH) using two different probes against the *Rbfox3* coding sequence and its 3’UTR. Consistent with the scRNA-Seq results, *Rbfox3* ISH signals were enriched in the SVZ/IZ(subventricular zone/intermediate zone) at E14.5 and partially co-localized with Eomes-positive cells (Fig. 3d-3e and S3c). In contrast, *Rbfox1* and *Rbfox2* RNA ISH showed peak signals in the cortical plate (neurons), with *Rbfox2* also showing expression in the SVZ/IZ (Fig. 3d, adapted from Genepaint(*48*)). In E14.5 dorsal forebrains, Rbfox1/2/3 showed overlapping protein expression in the cortical plate, with Rbfox3 and Rbfox2 emerging in the SVZ/IZ (Fig. 3f-3g). These results suggest that *Rbfox3/2* are expressed in immature neurons and all three Rbfox genes are expressed in the cortical plate at E14.5.

Analyses of a scRNA-Seq dataset(*47*) spanning E10.5-P4 mouse cortical cells showed that *Rbfox1/2/3* genes displayed variable expression levels among cortical neuron subtypes, with *Rbfox3* showing higher expression in Layer IV-V, VIb, and other deep layer neurons (Fig. S3d-S3e). Immunostaining of P0 neocortices confirmed that Rbfox3 displayed higher expression in layer IV-V and subplate neurons, while Rbfox1 showed higher expression in layer V-VI (Fig. 3h and S3f). These results suggest that *Rbfox* genes display overlapping and variable expression among cortical layers at P0.

### Ectopic Rbfox expression in RGCs increases neuronal splice isoforms and promotes neuronal differentiation

We introduced *Rbfox3* into RGCs at E13.5 by in utero electroporation (IUE) and analyzed cell states and positions at E14.5 (Fig. S4a-S4c) and E15.5 (Fig. S4d-S4f). 24 hours after IUE, Rbfox3 expression did not significantly change the overall cell distribution, or the proportions of Pax6- or Eomes-positive cells (Fig. S4a-S4c). Two days after IUE, the ectopic Rbfox3 significantly decreased the proportion of cells in the VZ (Fig. S4d-S4e), as well as the proportions of Pax6-positive RGCs and Ki67-positive cells (Fig. S4d and S4f). Pulse BrdU labeling confirmed the reduction of proliferating cells in the VZ (Fig. S4d and S4f). These observations indicate that ectopic Rbfox3 promotes RGC-to-neuron differentiation.

To identify molecular targets of Rbfox3, we performed IUE at E13.5, isolated the cells with FACS at E14.5, and performed RNA-Seq (Fig. 4a). Transient *Rbfox3* expression significantly altered the splicing profiles in RGCs, including 1694 SEs (Fig. 4b). Gene Ontology analysis of differentially spliced genes uncovered that ATP-binding proteins, zinc-finger, and transcription regulators were most highly enriched terms (Fig. 4c). The canonical Rbfox binding motif *GCAUG* was highly enriched in the downstream introns of up-regulated exons, and the same motif was highly enriched inside SEs that showed decreased inclusion upon *Rbfox3* expression (Fig. 4d). Expression of *Rbfox3* alone regulated 334 neuron-progenitor differentially spliced exons, with 160 exons upregulated in neurons and 50 exons downregulated; importantly, *Rbfox3* transient expression regulated 50 out of 136 transcription regulators that showed higher inclusion in neurons and promoted the inclusion for 21 of them (Fig. 4e-4g, Fig. S4g-S4h). These numbers are probably underestimated because of the stringent threshold (|ΔPSI| > 0.15 and FDR < 0.05). Comparably, transient expression of Rbfox2 or Rbfox1 in RGCs also redirected the splicing of transcription regulators toward neuronal isoforms (Fig. 4h and Fig. S4i). These results support that *Rbfox* expression directs hundreds of genes, especially transcription regulators, to their neuronal isoforms.

**Figure 4.**
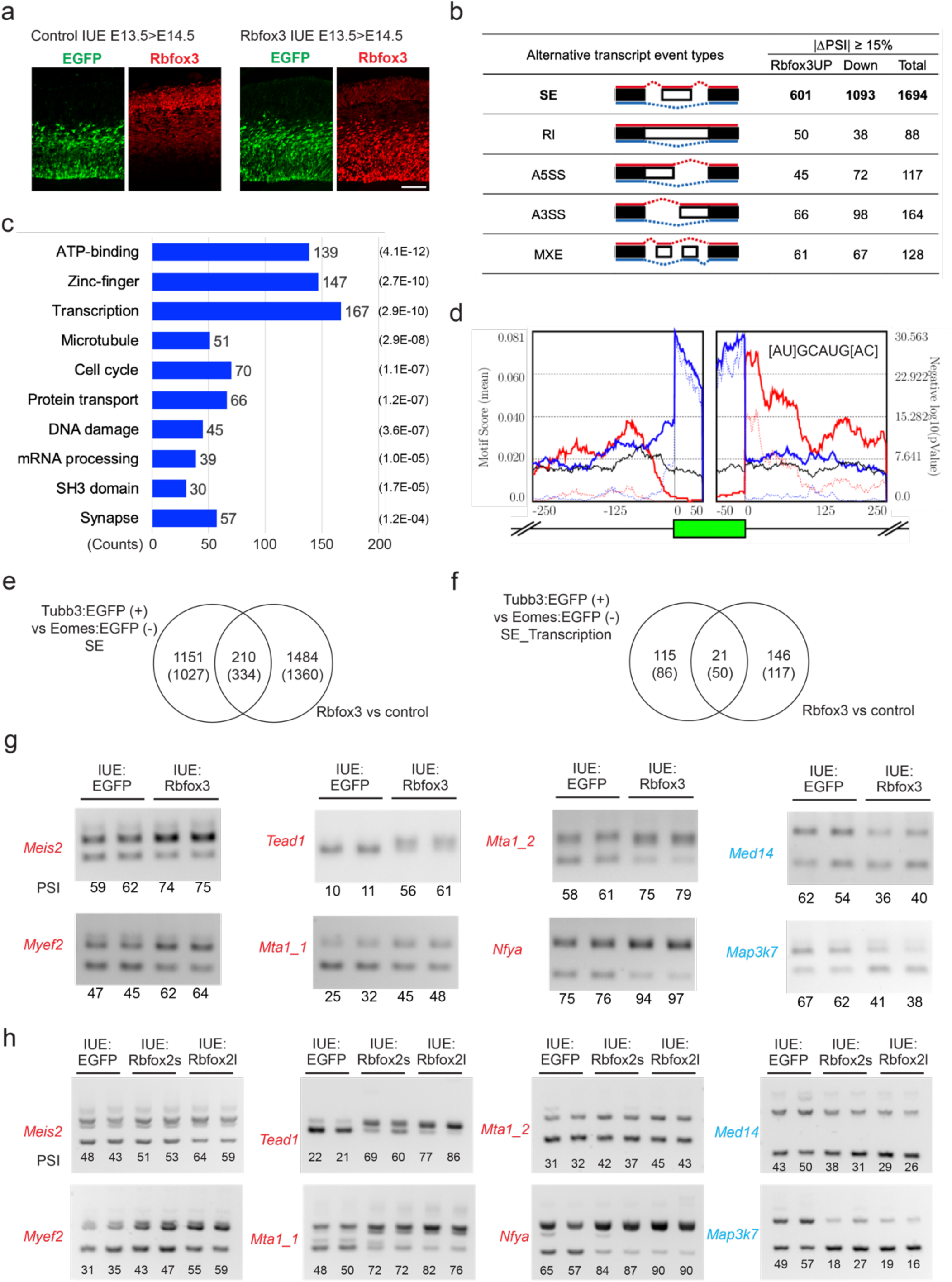
Rbfox proteins promote transcription regulators to neuronal splice isoforms. **a.** Transient expression of Rbfox3 in the radial glial progenitors by IUE at E13.5 and stained on E14.5. Green, antibody against EGFP which was expressed by both the control and Rbfox3 vectors (pCAG-ires-EGFP); red, anti-Rbfox3 showing both endogenous and ectopic Rbfox3; scale bar, 100μm. **b.** RNA-Seq analysis of FACS-sorted Rbfox3-ires-EGFP positive cells showing extensive splicing changes when compared with EGFP control cells (|ΔPSI| >= 15%, FDR < 0.05). **c.** Gene ontology analysis showing that transcription regulators were highly enriched for Rbfox3-mediated alternative splicing. Top-to-bottom ranked by enrichment score. The FDR (numbers in parentheses) and number of affected gene counts are indicated. **d.** The *GCAUG* motif was highly enriched in the downstream introns of Rbfox3 up-regulated exons. The solid red and blue lines show the motif enrichment score in up- and down-regulated exons; the dashed red and blue lines show the statistical significance of motif enrichment in up- and down-regulated exons, respectively. The green box shows the alternative exon, and integers indicate relative base positions to the beginning or end of the alternative exon. **e.** Transient expression of Rbfox3 (IUE E13.5 > E14.5) promoted the inclusion of 160 neuron-enriched exons (Tubb3:EGFP(+) cells) and the skipping of 50 progenitor-enriched exons (Eomes:EGFP(-) cells, p = 7.38e-07 by Hypergeometric test). 334/1361 exons responded to the Rbfox3 expression. |ΔPSI| < 0.15 and FDR < 0.05. **f.** Transient Rbfox3 expression (IUE E13.5 > E14.5) up-regulated 21 neuron-enriched exons in transcription regulators (p = 2.17e-18 by Hypergeometric test). 50/136 exons in transcription regulators responded to Rbfox3. **g.** RT-PCR validation of the effects of transient Rbfox3 expression (IUE on E13.5 > analyses on E14.5) on neuron-specific exons such as in *Meis2* and *Tead1*, and RGC-specific exons in *Map3k7* and *Med14*, numbers indicate PSI values. Red and blue colors of gene names indicate up- or down-regulation of exon inclusion upon Rbfox3 expression, respectively. **h.** RT-PCR validation of the effects of Rbfox2 isoforms on neuron-specific exons (IUE E13.5 > E14.5). See also Figure S4.

### *Rbfox1/2/3* triple CRISPR knockout impairs radial neuronal migration

Rbfox1/2/3 proteins show similar protein structures, overlapping expression in postmitotic neurons (Fig. 3 and S3), and genetic compensation in mice (*30, 31*). To understand their redundant function, we expressed three guide RNAs targeting each of *Rbfox1/2/3* in one construct (*tKO*) and electroporated it into the developing mouse dorsal forebrain at E13.5. We collected brain sections at P0 and used immunostaining of Rbfox1/2/3 separately and in combination to evaluate the knockout efficiency. In wild-type, 87.9% of Cas9-2A-EGFP positive cells (control) expressed at least one of the three Rbfox1/2/3 proteins, while in *Rbfox1/2/3 tKO*, 73.2% of Cas9-2A-EGFP-triple-gRNA cells were negative for all three Rbfox1/2/3 proteins by immuno-staining (Fig. 5a-5b). We quantified the position of *Rbfox1/2/3 tKO* cells at P0 and found excess cells in deep layers and reduced cell numbers in the superficial layers (Fig. 5c). We co-stained upper layer marker Cux1 and deep layer marker Ctip2 and found that Rbfox1/2/3 did not significantly change Cux1 or Ctip2 proportion (Fig. S5a-S5b).

**Figure 5.**
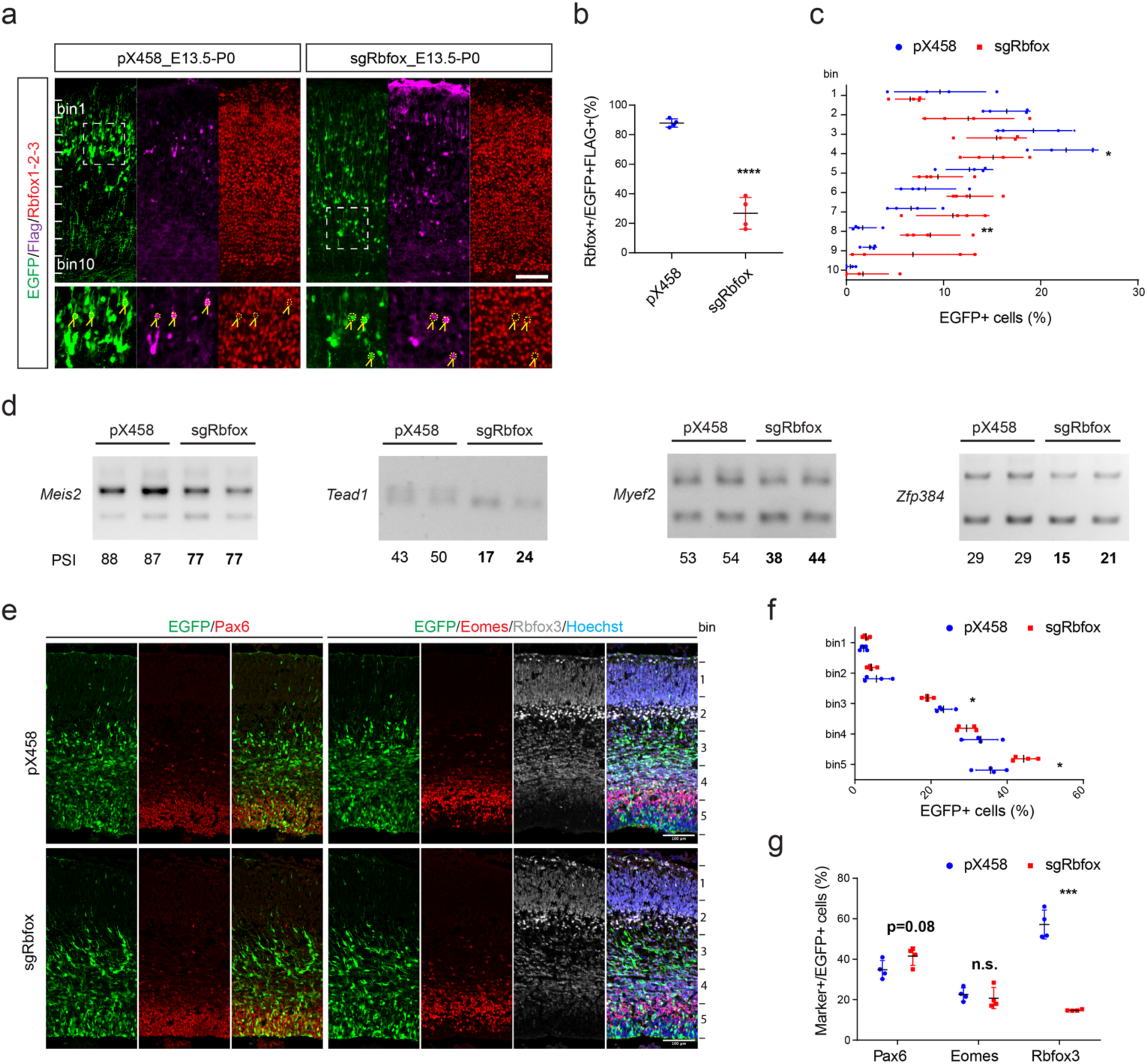
Simultaneous ablation of Rbfox1/2/3 impairs radial neuronal migration. **a.** Simultaneous *in-utero* delivery of *CRISPR/Cas9* guide RNAs at E13.5 significantly decreased Rbfox1/2/3 expression at P0 (antibodies to all three Rbfox1/2/3 proteins, red) and led to neuronal migration defects. EGFP (green) shows 2A-EGFP in the construct, and Flag (purple) shows anti-Flag signals where the Flag tag was fused with SpCas9. Scale bar, 100μm. **b.** Quantification of Rbfox1/2/3 positive cells (mixture of three Rbfox1/2/3 antibodies) in control and *Rbfox1/2/3* triple knockout brain slices (unpaired two-tailed Student’s t-test, n=4 sections from four P0 brains for each group). **c.** Quantification of Rbfox1/2/3 triple knockout (tKO) cells showing abnormal accumulation of cells in the deep layers and depletion of cells in the superficial layers (unpaired two-tailed Student’s t-test, n=4 P0 brains for each group). **d.** RT-PCR results showing that Rbfox1/2/3 triple knockout in the brain regulated alternative splicing of transcription regulators such as *Meis2* and *Tead1*. **e.** IUE of Rbfox1/2/3 tKO at E13.5 and examination of brain tissues at E15.5. Antibodies against Pax6 (red), Eomes (red), and Rbfox3 (grey). EGFP (green) shows 2A-EGFP in the CRISPR/Cas9 construct. Scale bar, 100μm. **f.** Quantification of the distribution of Rbfox1/2/3 tKO cells (EGFP+) in the embryonic neocortex (unpaired two-tailed Student’s t-test, n=4 sections from four E15.5 brains for each group). **g.** Portions of Pax6-, Eomes-, and Rbfox3-positive cells in EGFP-positive cells in control and *Rbfox1/2/3* triple knockout brain slices. Unpaired two-tailed Student’s t-test, n=4 sections from four E15.5 brains for each group. See also Figure S5.

We analyzed alternative splicing of *Meis2* exon12 in FACS sorted *Rbfox1/2/3 tKO* cells and found that its inclusion level was significantly decreased (10.8%, p < 0.001, t-test, Fig. 5d). The inclusion levels of neuron-specific exons were also decreased in *Rbfox1/2/3 tKO* cells for other transcription regulators such as *Tead1* and *Myef2* (Fig. 5d and S5c). We further analyzed the position of *Rbfox1/2/3 tKO* cells in E15.5 brains (IUE at E13.5) and found significantly more cells in the VZ (Fig. 5e-5f). Consistently, there were more Pax6- and EGFP-double-positive cells in the *Rbfox1/2/3 tKO* brains (Fig. 5g). In summary, these results indicate that *Rbfox1/2/3* proteins are required for radial neuronal migration and alternative splicing of transcription regulators in the developing neocortex.

### Rbfox proteins promote *Ptbp1* mammal-specific alternative exon8 and a poison exon inclusion

*Ptbp1* suppresses neuronal exon inclusion (*3, 13, 38*), and the mouse *Ptbp1* exon8 (human *PTBP1* exon9) has been reported to cause evolution differences between vertebrates(*39, 40*). Unbiased analysis of *Rbfox3* upregulated exons uncovered *Ptbp1* exon8 (Fig. 6a-6b). We identified a conserved *GCAUG* Rbfox binding motif downstream of *Ptbp1* exon8 (motif_1, Fig. 6b and S6a) and validated that transient *Rbfox3* or *Rbfox2* expression in cortical RGCs promotes *Ptbp1* exon8 inclusion (Fig. 6c and S6b). *Rbfox1/2/3 tKO* significantly decreased *Ptbp1* exon8 inclusion in Neuro2a cells (Fig. S6c). These results indicate that Rbfox proteins promote *Ptbp1* exon8 inclusion.

**Figure 6.**
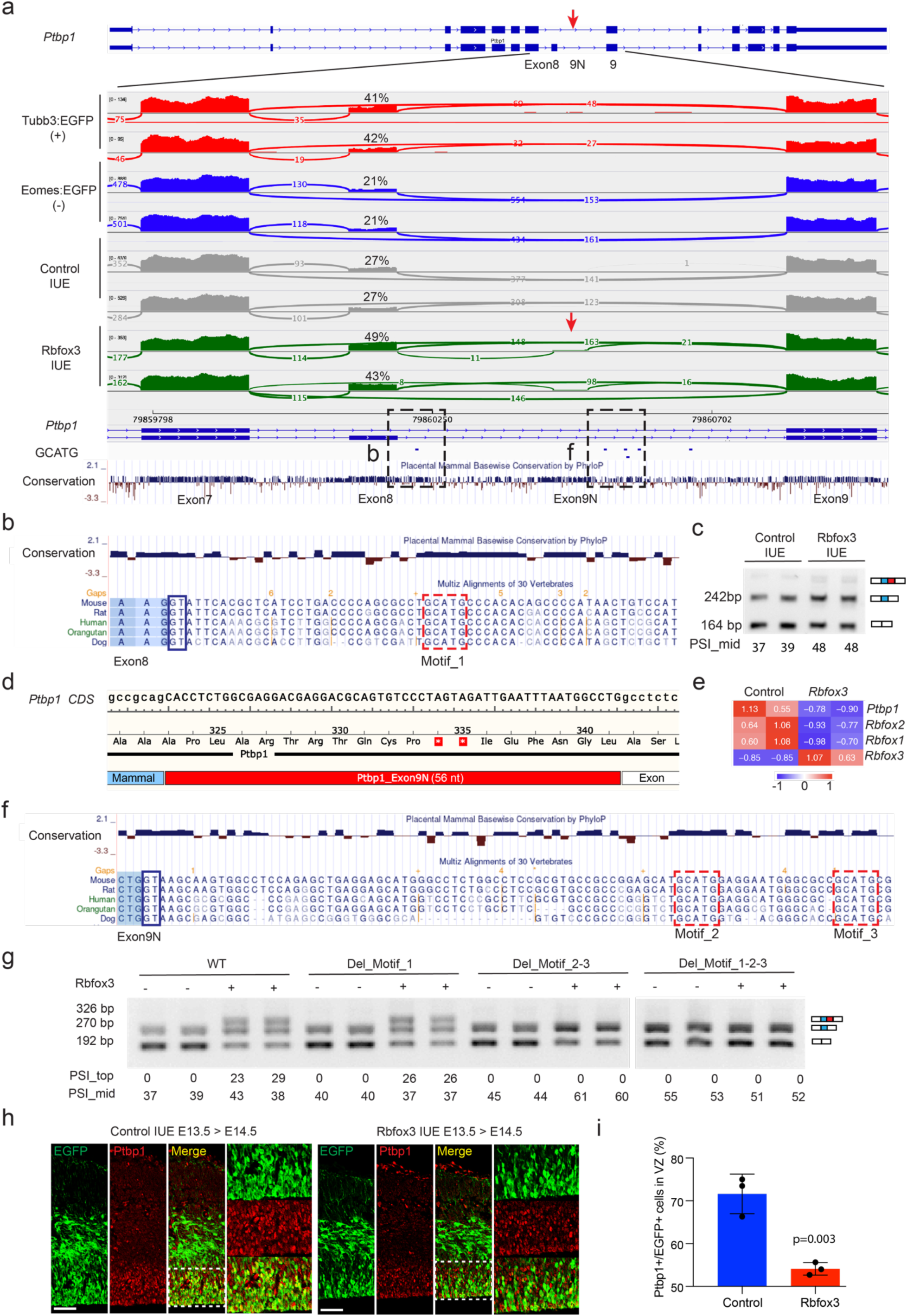
Rbfox3 promotes the *Ptbp1* mammal-specific exon8 and poison exon9N inclusion. **a.** Sashimi plots showing that *Ptbp1* mammal-specific alternative exon8 (B) displayed higher inclusion in cells enriched for neurons (*Tubb3:EGFP*(+)) than RGCs (*Eomes:EGFP*(-)), and transient expression of Rbfox3 (IUE) increased its inclusion levels *in vivo*. In addition, *Ptbp1* poison exon9N (Fig. 6d below, red arrow) was upregulated by Rbfox3 expression. Conservation scores at the bottom show that the two alternative exons and the *GCAUG* motifs are conserved in mammals. **b.** Zoom in view of the downstream intron of *Ptbp1* exon8 showing the conserved *GCAUG* Rbfox binding motif_1. **c.** RT-PCR results show that transient Rbfox3 expression in RGCs promoted *Ptbp1* exon8 inclusion. **d.** *Ptbp1* poison exon9N introduces premature stop codons that were predicted to induce NMD. **e.** Heat map of RNA-Seq results showing that transient Rbfox3 expression downregulated *Ptbp1*, *Rbfox1*, and *Rbfox2* mRNA levels. Colors indicate scaled expression. **f.** The downstream intron of Ptbp1 poison exon9N contains two conserved *GCAUG* Rbfox binding motif_2 and motif_3. **g.** Mini gene-splicing assays in Neuro2a cells showing that Rbfox3 expression promoted *Ptbp1* exon8 and exon9N inclusion. Deleting Motif_1 attenuated the Rbfox3-mediated effect on exon8, while Motifs_2/3 were required for Rbfox3-mediated exon 9N inclusion. Numbers indicate PSI values. **h.** Transient Rbfox3 expression by IUE (E13.5 > E14.5) leads to a decreased proportion of Ptbp1-positive cells (yellow/green). Green, anti-EGFP; red, anti-Ptbp1. Scale bar, 50μm. **i.** Quantifications of Ptbp1-positive cells in the VZ of control and Rbfox3 IUE brain slices (unpaired two-tailed Student’s t-test, n=3 sections from two E14.5 brains for each group). See also Figure S6.

Very interestingly, expression of Rbfox3 led to the inclusion of a conserved 56-nt poison exon9N in *Ptbp1* and significantly decreased *Ptbp1* mRNA levels (decrease by 31%, adj.p < 0.001, Fig. 6a, 6d-6e). Rbfox3 expression also led to decreased *Rbfox1* and *Rbfox2* mRNA levels likely through splicing (Fig. 6e and S6d-S6e). We identified two conserved *GCAUG* motifs (motif_2 and motif_3) in the downstream intron of *Ptbp1* exon9N (Fig. 6f and S6a). We created a *Ptbp1* minigene (spanning exon6-9) and found that *Rbfox3* or *Rbfox2* expression promoted the inclusion of *Ptbp1* exon8 and the poison exon9N (Fig. 6g and S6f). We further mutated the *GCAUG* motifs and found that motif_1 and motifs_2_3 were required for *Rbfox3/2* mediated *Ptbp1* exon8 and exon9N inclusion, respectively (Fig. 6g). Immunostaining of Rbfox3 IUE brain slices showed that the proportions of Ptbp1-positive cells in the VZ were significantly decreased upon Rbfox3 expression (72% control versus 54% Rbfox3, n=3, Fig. 6h-6i). These results support that Rbfox proteins promote *Ptbp1* mammal-specific alternative exon8 and antagonize *Ptbp1* expression by directly promoting poison exon9N inclusion.

### Meis2 splice isoforms differentially regulate neurogenesis and neuronal migration

To understand the regulation of transcription regulator isoforms, we focus on the *Meis2* gene, which displays differential splicing of exon12 between RGCs (exon12 skipped) and neurons (exon12 included, Fig. 1f-1g). Expression of *Rbfox3* promoted *Meis2* exon12 inclusion (Fig. 4g), and re-analysis of CLIP-Seq results from an independent study(*33*) showed that *Rbfox3* bound directly to *Meis2* exon12 (Fig. 7a). Close examination of *Meis2* exon12 uncovered two consecutive and conserved *GCAUG* Rbfox binding motifs (Fig. 7b). We mutated the two *GCAUG* sites to *G<u>U</u>AUG* in a mini gene construct and found that the inclusion levels of *Meis2* exon12 were significantly decreased by 43% (Ex12-mut, Fig. 7c). Rbfox2 (Tag1, Del_1) and Rbfox1 (Tag2 and Tag3, Del_2 and Del_3) showed CLIP tags in the upstream and downstream introns, respectively (Fig. 7a). We deleted these binding motifs individually or together with exon12 mutations in the minigene constructs and examined their effects on splicing (Fig. 7c). Introduction of Del_1, Del_2, or Del_3 alone (deletion of CLIP Tag1-3 sequence, Fig. 7a) had small or no effect, while these deletions showed moderate synergistic effects when combined with exon12 mutations (Fig. 7c). These results indicate that Rbfox proteins promote *Meis2* exon12 inclusion predominantly by binding to *GCAUG* motifs inside exon12.

**Figure 7.**
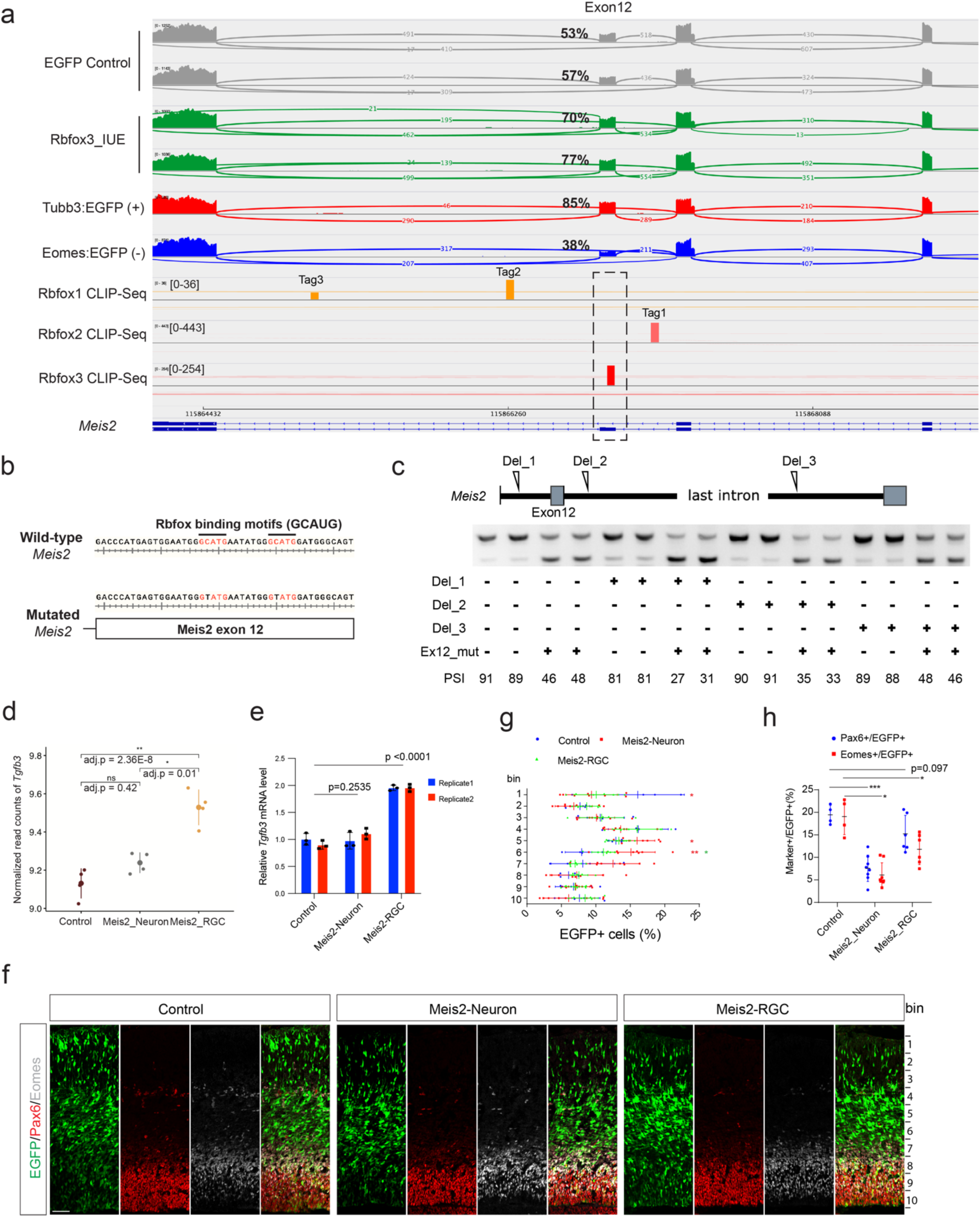
Rbfox-mediated splicing of Meis2 leads to functionally different isoforms. **a.** Sashimi plots showing the alternative splicing of *Meis2* exon12 upon Rbfox3 expression (IUE E13.5 > E14.5) and Rbfox3/2/1 Clip-Seq tags (dashed rectangle for Rbfox3, Tag1 for Rbfox2, Tag2 and Tag3 for Rbfox1). The Clip-Seq results were derived from a previous study(*71*). **b.** *Meis2* minigene constructs showing that the two Rbfox binding motifs (*GCAUG*) in *Meis2* exon12 were mutated to *G<u>U</u>AUG*. **c.** Mini gene-splicing assay results show that mutating Rbfox binding motifs within *Meis2* exon12 (Fig. 7b) substantially inhibited its inclusion. Del_1, as well as Del_2 and Del_3 (Deletion of Tag2 and Tag3), showed synergistic effects with exon12 mutations. **d.** RNA-Seq results (variance stabilizing transformed read count, with standard deviation) showing that ectopic expression of *Meis2* RGC-isoform (exon12 skipped), but not the neuron-isoform, promoted *Tgfb3* mRNA expression. Differential gene expression was determined with the Wald test, corrected by the Benjamini and Hochberg method, and adjusted p-values are shown. **e.** RT-PCR validation of differential *Tgfb3* expression regulated by *Meis2* isoforms (bars showing SD, one-way ANOVA followed by Dunnett’s multiple comparisons test). **f.** Transient expression of empty vector or *Meis2* isoforms in the radial glial progenitors by IUE at E13.5 and examining brain tissues at E15.5. Antibodies against EGFP (green), Pax6 (red), and Eomes (grey). Scale bar, 100μm. **g.** Quantification of the distribution of EGFP-positive cells in the embryonic neocortex. Unpaired two-tailed Student’s t-test, n=4 sections from four E15.5 brains for the control group, n=8 sections from five brains for the *Meis2* neuron isoform group, and n=6 sections from three brains for the *Meis2* RGC group. **h.** Quantification of Pax6- and Eomes-positive cells in EGFP-positive cells in control and *Meis2* neuron- or RGC-isoform brain slices. Unpaired two-tailed Student’s t-test, n=4 sections from four E15.5 brains for the control group, n=8 sections from five brains for the *Meis2* neuron isoform group, and n=6 sections from three brains for the *Meis2* RGC group. See also Figure S7.

We further compared the functions of *Meis2* isoforms in gene regulation and neurogenesis. First, we expressed *Meis2* RGC isoform (skipping exon12) and neuronal isoform (exon12 included) in Neuro2a cells and performed RNA-Seq analyses with 4 replicates for each isoform and the empty-vector control (Fig. S7a). The two Meis2 isoforms were expressed at comparable levels (statistically not significant, Fig. S7b). We identified several differentially expressed genes between the two *Meis2* isoform sample groups, and of particular interest was the *Tgfb3* gene which was upregulated by the *Meis2* RGC isoform but not the neuronal isoform (Fig. 7d). Differential *Tgfb3* expression was cross validated by qPCR (Fig. 7e). *Tgfb3* encodes a TGF-β ligand and has been reported to promote cortical neurogenesis together with *Tgfb2*(*49*). Consistent with the *Meis2* RGC isoform expression, *Tgfb3* transcripts were enriched in the ventricular zone (VZ, Fig. S7c, adapted from Genepaint(*48*)). When electroporated into the mouse brain at E13.5 and examined at E15.5, the *Meis2* neuronal isoform significantly stalled cells in the intermediate zone (bins 5-6) and decreased neurons in the cortical plate (bin 1, Fig. 7f-7g). The *Meis2* neuronal isoform decreased proportions of Pax6- or Eome-positive cells, while the *Meis2* RGC isoform showed a significantly milder effect (Fig. 7h). These results suggest different functions of *Meis2* isoforms in regulating gene expression and cortical neurogenesis.

## Discussion

The cell-type-specific splicing analysis in this study uncovered an enrichment of transcription regulators during cortical development. This observation was further supported by isoform discovery from single-cell long-read sequencing. Over one dozen alternative splicing events were predicted to trigger NMD in transcription regulators and another 23 events changed protein domain structures. Our results indicate that alternative splicing events, especially the ones mediated by Rbfox proteins, directly rewire transcription regulators during cortical neurogenesis, providing mechanistic insights how RBFOX1 knockdown affected gene transcription(*50*).

*Ptbp1* has been reported to suppress neuronal fate *in vitro* (*51*) and *Ptbp1* knockout causes lethal hydrocephalus in mice (*34*). The mammal-specific *Ptbp1* alternative exon8 has been reported to evolve from a constitutive exon in lower organisms(*39*) and alter splicing profiles among vertebrates(*39, 40*). Here, we show that Rbfox3/2 proteins promote the inclusion of *Ptbp1* exon8. While Ptbp1 and Rbfox proteins have been reported to regulate neuronal exons synergistically(*3, 13*), whether they crosstalk directly remained unknown. We show that Rbfox proteins suppress Ptbp1 by directly promoting the insertion of exon9N which is predicted to introduce premature stop codons and cause AS-NMD. These results suggest that Rbfox3/2 directly regulate *Ptbp1* by promoting the poison exon insertion.

The Meis2 RGC and neuronal isoforms differentially regulate *Tgfb3* expression and cortical neurogenesis. Recently, the Rbfox-regulated *Tead1* exon has been shown to regulate its transcription activity in HEK293 cells(*52*). Thus, Rbfox3/2 proteins antagonize *Ptbp1* and differentially splice transcription factors such as *Meis2* and *Tead1* to promote neuronal differentiation. Meis2 and Pbx1 form a protein complex for transcription activation(*53*). Previous studies of *Pbx1*(*54, 55*) and our present results of *Meis2* showed that *Pbx1* and *Meis2* are alternatively spliced by Ptbp and Rbfox proteins, respectively. These observations suggest that splicing factors can synergistically regulate transcription co-regulators.

Members of RNA splicing regulator families frequently show overlapping expression in the brain, presenting technical challenges to fully understanding their redundant functions. Gene compensation and neurological deficits have been reported in *Rbfox1/2/3* single knockout mice: *Rbfox1* knockout mice have been reported to show normal brain morphology with increased susceptibility to seizures(*31*), and *Rbfox1* knockdown has been reported to affect neuronal migration and dendritic arborization(*56*); *Rbfox2* deletion in the central nervous system leads to a smaller cerebellum with ectopic localization of Purkinje cells(*30*); *Rbfox3* knockout has been reported to cause reduced brain weight and increased seizure vulnerability(*57*). In contrast, *Rbfox1/2/3* proteins have been reported to regulate thousands of splicing events and triple *Rbfox1/2/3* knockout has been reported to impair axon initial segment assembly in motor neurons derived from mouse embryonic stem cells(*58*). Here, simultaneously knocking out *Rbfox1/2/3* genes in the developing neocortex with CRISPR/Cas9 led to defects in radial neuronal migration and dysregulated splicing patterns. These results suggest that multiplexed CRISPR editing is robust in understanding redundant gene functions in the developing neocortex. The direct molecular cause of neuronal migration defects following *Rbfox1/2/3* ablation remains unclear, which is conceivably the combinatory outcome of splicing changes, such as the ones in cytoskeletal proteins and transcription regulators, and the non-splicing effects on mRNA modification and protein translation(*13, 50, 59, 60*).

## Methods and Materials

### Mouse protocols

Mouse protocols were reviewed and approved by University of Chicago Institutional Animal Care and Use Committee. The *Tg(Eomes/Tbr2:EGFP)* and *Tg(Tubb3:EGFP)* Bac transgenic mouse lines, for which *EGFP-PolyA* cassettes were inserted in front of the translational stop codons of *Eomes* and *Tubb3*, were obtained from GENSAT(*41*) and crossed to the FVB inbreed strain for at least two generations. Heterozygous transgenic mice did not display any obvious developmental defects. To isolate cells, the dorsal forebrains were dissected from E14.5 heterozygous *Tg(Tbr2-EGFP)* or *Tg(Tubb3:EGFP)* transgenic mice, dissociated with the Papain (Worthington) and resuspended in presort medium (Neural Basal medium supplemented with 2% B27, 1% N2, 1% Penicillin-streptomycin, 10ng/ml EGF, 10ng/ml FGF2, 0.5% FBS and 0.25% HEPES). Single-cell suspension was cooled on ice and immediately sorted on the FACSAria II sorter (BD). EGFP-positive and -negative populations were collected into post-sort medium (same as presort medium but with 20% FBS) and spun down for subsequent analyses.

### RNA-Sequencing and data analysis

Total RNA was extracted from FACS-sorted primary cells and cell lines, subjected to stranded mRNA-Seq library preparation following standard protocols (Illumina Tru-Seq) and paired-end sequencing on Illumina Nextseq500 and Novaseq6000 platforms at the University of Chicago. Three to five IUE-positive embryonic dorsal cortices of the same litter or batch were pooled together before Papain dissociation (Worthington), and about 100,000 cells were sorted directly into Trizol for RNA extraction with Direct-zol (Zymo). 100 ng total RNA was used as input for Tru-Seq library prep (Illumina). Raw reads were trimmed and aligned to mouse mm10 with STAR aligner(*61*). Alternative splicing was analyzed with rMATS(*62*), enriched motif analysis was done with rMAPS(*63*), and gene ontology was analyzed with DAVID(*64*). Differential gene expression was analyzed with Salmon(*65*) and DESeq2(*66*). Rbfox1/2/3 CLIP-Seq results(*33*) were aligned to the mouse genome and significant peaks were called by CTK(*67*). Protein domains were predicted using the NCBI Conserved Domain Database, compared between pairs of splice isoform sequences, and located to SE events with customized scripts. Protein 3-D structures were modeled by SWISS-MODEL and presented by open-source PyMOL v2.5.0. In parallel to our scRNA-Seq analysis of E14.5 CD1 dorsal cortex previously reported(*44*), the untagmented full-length cDNA was further amplified and used for long-read sequencing with PacBio Sequel II. PacBio library was prepared with 1μg amplified cDNA using SMRTbell Express Template Prep Kit V2.0. The differential splicing between pooled long reads was analyzed using FLAIR(*45*).

### Molecular cloning, cell culture and RT-PCR

The *Meis2* mini gene construct (pR085): genomic DNA of Neuro2a cells was amplified with primer pairs XR301-XR302 and inserted to pCAG-HF-IRES-EGFP by Gibson Assembly (NEB).

*Meis2* mutated mini gene (pR086): the mutations in the Rbfox3 binding motifs were created by amplifying genomic DNA of Neuro2a cells with primer pairs XR301-XR304 and XR303-XR302 and cloned into pCAG-HF-IRES-EGFP by Gibson Assembly (NEB).

The *Meis2* neuron isoform (NM_001159567) expression construct (pR084-v3): E14.5 mouse brain complementary DNA (cDNA) was amplified with primers pairs XR293-XR294 and cloned into pCAG-HF-IRES-EGFP by Gibson Assembly (NEB).

The *Meis2* RGC isoform (NM_001159568) expression construct (pR084-v4): E14.5 mouse brain cDNA amplified with primers pairs XR293-XR296 and cloned into pCAG-HF-IRES-EGFP by Gibson Assembly (NEB).

The *Ptbp1* wild-type mini-gene construct (pZ046): genomic DNA of C57BL/6J was amplified with primer pairs CH986-CH987 and ligated to Asc1-Not1 linearized vector pCAG-HF-ires-Puro.

The *Ptbp1* motif_1 deletion construct (pZ047): genomic DNA of C57BL/6J was amplified with primer pairs CH986-CH990 and CH991-CH987, and the two fragments inserted into Asc1-Not1 linearized pZ046 by Gibson Assembly (NEB).

The *Ptbp1* motif_2-3 deletion construct (pZ048): C57BL/6J genomic DNA was amplified with primer pairs CH986-CH992 and CH993-CH987, and the two fragments were inserted into Asc1-Not1 linearized pZ046 by Gibson Assembly (NEB).

The *Ptbp1* motif_1-2-3 deletion construct (pZ048T): pZ047 (motif_1 deletion) was amplified with primer pairs CH986-CH992 and CH993-CH987, and the two fragments were inserted into Asc1-Not1 linearized pZ046 by Gibson Assembly (NEB).

The *Rbfox1/2/3* triple CRISPR knockout construct (pX458-Rbfox1-2-3-gRNA): three guide RNA cassettes CH687 gBlock (*Rbfox1*), CH690-CH691 (*Rbfox2*), and CH692-CH693(*Rbfox3*) were inserted into PciI and KpnI linearized pX458.

Plasmids described above or in our previous work(*13*) were transfected into Neuro2a cells (ATCC) with Fugene 6 (Promega). The transfected cells were selected by puromycin and subjected to RNA extraction with Trizol (Sigma). Primary hippocampal neurons were isolated from E18.5 CD1 mouse embryos with Papain (Wathington) and cultured for one day *in vitro*. Primary neurons were treated with DMSO or cycloheximide (100 μg/ml) for 5 hours before RNA extraction. Reverse transcription was performed with random primers following the manufacturer’s protocols (Superscript IV, Thermo Scientific). *Meis2* alternative exon was amplified with primer pairs XR244-XR245; *Ptbp1* alternative exons were amplified with primer pairs CH438-CH989 in Fig. 6g, and with CH988-CH989 (mouse *Ptbp1*) in Fig. 6c and Fig. S6f. *Tgfb3* qPCR was performed with primer pairs XR902-XR903. Primer sequences are listed in Table S1.

### In utero electroporation

CD-1 mice were purchased from Charles River, housed at the University of Chicago Animal Care Facility. At noon on the day when a vaginal plug was found was designated E0.5. The day of birth was designated postnatal day 0 (P0). IUE procedures were performed in accordance with previously described protocols(*68*). Briefly, the proper concentration (1-2 μg/μl) of DNA solution with 0.05% Fast Green dye was injected into the E13.5 brain lateral ventricle *in utero* via glass micropipettes, and 5 square fixed-potential pulses with a duration of 35-50 ms per pulse were administered by an Electro Square Porator (ECM830, BTX).

### Immunostaining of brain sections

Immunostaining was done as previously described(*44*). Briefly, embryonic brains were fixed in 4% paraformaldehyde (PFA) overnight at 4C°, cryoprotected in 25% sucrose overnight at 4C°, embedded in Frozen Section Medium (Thermo Scientific, Cat. 6502), and sectioned at 14μm in the coronal direction if not specified otherwise. Slices were rinsed with 1x PBS for 5min, incubated with blocking buffer (1x PBS containing 0.03% Triton X-100 and 5% normal donkey serum) at room temperature for 30 mins, and further incubated with primary antibodies diluted in PBST buffer (1x PBS containing 0.03% Triton X-100) overnight at 4C°. After 3 times washing with 1x PBS, slides were incubated for one hour at room temperature with fluorophore-conjugated secondary antibodies in the dark. Slides were scanned with a Leica SP5 confocal or Zeiss Apotome 2 microscope. The following primary antibodies were used: anti-Rbfox1 (Millipore MABE159, mouse, 1:800; Sigma-Aldrich HPA040809, rabbit, 1:500), anti-Rbfox2 (Bethyl A300-864A, rabbit, 1:500), anti-Rbfox3/NeuN (Cell Signaling Technology 24307T, rabbit, 1:500; Millipore MAB377, mouse, 1:500), anti-Cux1(Santa Cruz sc-13024, rabbit, 1:1000), anti-Ctip2 (Abcam ab18465, rat, 1:1000), anti-Tuj1 (BioLegend 801201, mouse, 1:1000), anti-GFP (Abcam ab13970, chicken, 1:3000), anti-Flag (Sigma-Aldrich F1804, mouse, 1:500), anti-Ptbp1 (Abcam ab5642, goat, 1:200), anti-Tbr2 (Thermo Fisher Scientific 14-4875-82, rat, 1:200), anti-BrdU (AbD Serotec MCA2060GA, rat, 1:500), anti-Ki67 (Abcam, ab15580, 1:200). Hoechst 33342 (Thermo Scientific, 62249) and the secondary antibodies were diluted at 1:2000 in PBST buffer: Donkey anti-chicken 488 (Jackson ImmunoReseach, 703-546-155), donkey anti-mouse 594 (Thermo Scientific, A21203), donkey anti-mouse 488 (Thermo Scientific, A21202), donkey anti-rabbit 488 (Thermo Scientific, A21206), donkey anti-rabbit 594 (Thermo Scientific, A21207), donkey anti-rat 647 (Thermo Scientific, A48272) and donkey anti-goat 647 (Thermo Scientific, A21447) and donkey anti-goat 594 (Thermo Scientific, A11058).

### In situ hybridization

The in situ hybridization (ISH) protocol was described previously(*69*). Digoxigenin-labeled antisense RNA probes were *in vitro* transcribed from PCR products amplified from the *Rbfox3* coding sequence (XR1038 and XR1039) and 3’UTR (XR1040 and XR1041). The co-immunostaining was performed following RNA ISH. Labeled slices were imaged using a Leica microscope.

### Statistical analyses

To identify differential splicing events, we included at least two biological replicates for each genotype, used built-in functions in rMATS, and filtered by False Discovery Rate (FDR) < 0.05 and ΔPSI indicated in individual figures. RT-PCR analyses of differential splicing (with two or more biological replicates) were compared with Student’s t-test assuming equal variance and with all details included in Table S2; Student’s t-test has been shown to be appropriate for extremely small sample sizes (*70*). p-values for scatter plots (Fig S1c and S4h) were calculated in R. Differential gene expression was determined with the Wald test, corrected by the Benjamini and Hochberg method, and adjusted p-values are shown (Fig. 7d and S7b). *Tgfb3* qPCR results (Fig. 7e) were compared using one-way ANOVA in GraphPad Prism, and means ± (standard deviation) were presented. Levels of significance were calculated with Hypergeometric Test for Fig. 7d-7e. In all figures if not specified otherwise: *, p-value < 0.05; **, p-value < 0.01; ***, p-value < 0.001; ****, p-value < 0.0001; n.s., p-value >0.05; bars indicate standard deviations.

## Data availability

Raw sequence data are available on NCBI SRA (PRJNA764749). All plasmids and codes are available upon request.

## Acknowledgment

The authors would like to thank all lab members for their valuable input and thank colleagues in the Department of Human Genetics, the Neuroscience Institute, and the DevNeuro group for their support. The long-read RNA-Seq was partially supported by the PacBio Iso-Seq SMRT Grant (2018). Works in the Zhang lab are supported by the NIGMS (DP2 GM137423, R35 GM152177) and the NIMH (R01 MH130594) to X.Z.

## Author contributions

X.R. and X.Z. led the project. X.R., K.H., Y.Y., R.Y., B.K., E.T., A.K., R.Z., and X.Z. performed experiments and analyzed data. K.H. led the RNA-Seq analyses and interpretation of exon functions. Y.Y. and E.T. led the analyses of long-read sequencing results. B.K. analyzed scRNA-Seq data. R.Y. assisted in RT-PCR validation of AS-NMD events. A.K. assisted in cell counting. R.Z. assisted in *Ptbp1* mini gene analysis. X.Z. supervised the project and wrote the manuscript with X.R. and other authors.

## References

1. E. T. Wang et al., Alternative isoform regulation in human tissue transcriptomes. Nature 456, 470–476 (2008).

2. Q. Pan, O. Shai, L. J. Lee, B. J. Frey, B. J. Blencowe, Deep surveying of alternative splicing complexity in the human transcriptome by high-throughput sequencing. Nat Genet 40, 1413–1415 (2008).

3. C. K. Vuong, D. L. Black, S. Zheng, The neurogenetics of alternative splicing. Nat Rev Neurosci 17, 265–281 (2016).

4. M. B. Johnson et al., Functional and evolutionary insights into human brain development through global transcriptome analysis. Neuron 62, 494–509 (2009).

5. H. Feng et al., Complexity and graded regulation of neuronal cell-type-specific alternative splicing revealed by single-cell RNA sequencing. Proc Natl Acad Sci U S A 118, (2021).

6. A. Joglekar et al., Single-cell long-read sequencing-based mapping reveals specialized splicing patterns in developing and adult mouse and human brain. Nat Neurosci, (2024).

7. A. A. Dillman et al., mRNA expression, splicing and editing in the embryonic and adult mouse cerebral cortex. Nat Neurosci 16, 499–506 (2013).

8. Y. Barash et al., Deciphering the splicing code. Nature 465, 53–59 (2010).

9. D. L. Black, Mechanisms of alternative pre-messenger RNA splicing. Annu Rev Biochem 72, 291–336 (2003).

10. E. L. Van Nostrand et al., A large-scale binding and functional map of human RNA-binding proteins. Nature 583, 711–719 (2020).

11. X. D. Fu, M. Ares, Jr., Context-dependent control of alternative splicing by RNA-binding proteins. Nat Rev Genet 15, 689–701 (2014).

12. D. Dominguez et al., Sequence, Structure, and Context Preferences of Human RNA Binding Proteins. Mol Cell 70, 854–867 e859 (2018).

13. X. Zhang et al., Cell-Type-Specific Alternative Splicing Governs Cell Fate in the Developing Cerebral Cortex. Cell 166, 1147–1162 e1115 (2016).

14. G. Panagiotakos et al., Aberrant calcium channel splicing drives defects in cortical differentiation in Timothy syndrome. Elife 8, (2019).

15. D. C. Lynch et al., Disrupted auto-regulation of the spliceosomal gene SNRPB causes cerebro-costo-mandibular syndrome. Nat Commun 5, 4483 (2014).

16. G. L. Carvill et al., Aberrant Inclusion of a Poison Exon Causes Dravet Syndrome and Related SCN1A-Associated Genetic Epilepsies. Am J Hum Genet 103, 1022–1029 (2018).

17. I. Voineagu et al., Transcriptomic analysis of autistic brain reveals convergent molecular pathology. Nature 474, 380–384 (2011).

18. S. De Rubeis et al., Synaptic, transcriptional and chromatin genes disrupted in autism. Nature 515, 209–215 (2014).

19. M. Irimia et al., A highly conserved program of neuronal microexons is misregulated in autistic brains. Cell 159, 1511–1523 (2014).

20. D. H. Geschwind, P. Rakic, Cortical evolution: judge the brain by its cover. Neuron 80, 633–647 (2013).

21. C. S. von Bartheld, J. Bahney, S. Herculano-Houzel, The search for true numbers of neurons and glial cells in the human brain: A review of 150 years of cell counting. J Comp Neurol 524, 3865–3895 (2016).

22. B. I. Bae, D. Jayaraman, C. A. Walsh, Genetic changes shaping the human brain. Dev Cell 32, 423–434 (2015).

23. J. H. Lui, D. V. Hansen, A. R. Kriegstein, Development and evolution of the human neocortex. Cell 146, 18–36 (2011).

24. M. Gotz, W. B. Huttner, The cell biology of neurogenesis. Nat Rev Mol Cell Biol 6, 777–788 (2005).

25. L. C. Greig, M. B. Woodworth, M. J. Galazo, H. Padmanabhan, J. D. Macklis, Molecular logic of neocortical projection neuron specification, development and diversity. Nat Rev Neurosci 14, 755–769 (2013).

26. R. F. Hevner, Intermediate progenitors and Tbr2 in cortical development. J Anat 235, 616–625 (2019).

27. K. Y. Kwan, N. Sestan, E. S. Anton, Transcriptional co-regulation of neuronal migration and laminar identity in the neocortex. Development 139, 1535–1546 (2012).

28. B. Raj et al., Cross-regulation between an alternative splicing activator and a transcription repressor controls neurogenesis. Mol Cell 43, 843–850 (2011).

29. J. Liu, A. Geng, X. Wu, R. J. Lin, Q. Lu, Alternative RNA Splicing Associated With Mammalian Neuronal Differentiation. Cereb Cortex 28, 2810–2816 (2018).

30. L. T. Gehman et al., The splicing regulator Rbfox2 is required for both cerebellar development and mature motor function. Genes Dev 26, 445–460 (2012).

31. L. T. Gehman et al., The splicing regulator Rbfox1 (A2BP1) controls neuronal excitation in the mammalian brain. Nat Genet 43, 706–711 (2011).

32. Y. S. Lin et al., Neuronal Splicing Regulator RBFOX3 (NeuN) Regulates Adult Hippocampal Neurogenesis and Synaptogenesis. PLoS One 11, e0164164 (2016).

33. S. M. Weyn-Vanhentenryck et al., HITS-CLIP and integrative modeling define the Rbfox splicing-regulatory network linked to brain development and autism. Cell Rep 6, 1139–1152 (2014).

34. T. Shibasaki et al., PTB deficiency causes the loss of adherens junctions in the dorsal telencephalon and leads to lethal hydrocephalus. Cereb Cortex 23, 1824–1835 (2013).

35. J. K. Vuong et al., PTBP1 and PTBP2 Serve Both Specific and Redundant Functions in Neuronal Pre-mRNA Splicing. Cell Rep 17, 2766–2775 (2016).

36. M. Zhang et al., Axonogenesis Is Coordinated by Neuron-Specific Alternative Splicing Programming and Splicing Regulator PTBP2. Neuron 101, 690–706 e610 (2019).

37. P. L. Boutz et al., A post-transcriptional regulatory switch in polypyrimidine tract-binding proteins reprograms alternative splicing in developing neurons. Genes Dev 21, 1636–1652 (2007).

38. Y. I. Li, L. Sanchez-Pulido, W. Haerty, C. P. Ponting, RBFOX and PTBP1 proteins regulate the alternative splicing of micro-exons in human brain transcripts. Genome Res 25, 1–13 (2015).

39. N. L. Barbosa-Morais et al., The evolutionary landscape of alternative splicing in vertebrate species. Science 338, 1587–1593 (2012).

40. S. Gueroussov et al., An alternative splicing event amplifies evolutionary differences between vertebrates. Science 349, 868–873 (2015).

41. S. Gong et al., A gene expression atlas of the central nervous system based on bacterial artificial chromosomes. Nature 425, 917–925 (2003).

42. J. C. Long, J. F. Caceres, The SR protein family of splicing factors: master regulators of gene expression. Biochem J 417, 15–27 (2009).

43. H. N. Jang et al., Exon 9 skipping of apoptotic caspase-2 pre-mRNA is promoted by SRSF3 through interaction with exon 8. Biochim Biophys Acta 1839, 25–32 (2014).

44. X. Ruan et al., Progenitor cell diversity in the developing mouse neocortex. Proc Natl Acad Sci U S A 118, (2021).

45. A. D. Tang et al., Full-length transcript characterization of SF3B1 mutation in chronic lymphocytic leukemia reveals downregulation of retained introns. Nat Commun 11, 1438 (2020).

46. N. Hug, D. Longman, J. F. Caceres, Mechanism and regulation of the nonsense-mediated decay pathway. Nucleic Acids Res 44, 1483–1495 (2016).

47. D. J. Di Bella et al., Molecular logic of cellular diversification in the mouse cerebral cortex. Nature 595, 554–559 (2021).

48. G. Diez-Roux et al., A high-resolution anatomical atlas of the transcriptome in the mouse embryo. PLoS Biol 9, e1000582 (2011).

49. T. Vogel, S. Ahrens, N. Buttner, K. Krieglstein, Transforming growth factor beta promotes neuronal cell fate of mouse cortical and hippocampal progenitors in vitro and in vivo: identification of Nedd9 as an essential signaling component. Cereb Cortex 20, 661–671 (2010).

50. B. L. Fogel et al., RBFOX1 regulates both splicing and transcriptional networks in human neuronal development. Hum Mol Genet 21, 4171–4186 (2012).

51. Y. Xue et al., Direct conversion of fibroblasts to neurons by reprogramming PTB-regulated microRNA circuits. Cell 152, 82–96 (2013).

52. S. Choi et al., RBFOX2-regulated TEAD1 alternative splicing plays a pivotal role in Hippo-YAP signaling. Nucleic Acids Res, (2022).

53. G. A. Bjerke, C. Hyman-Walsh, D. Wotton, Cooperative transcriptional activation by Klf4, Meis2, and Pbx1. Mol Cell Biol 31, 3723–3733 (2011).

54. A. J. Linares et al., The splicing regulator PTBP1 controls the activity of the transcription factor Pbx1 during neuronal differentiation. Elife 4, e09268 (2015).

55. L. Remesal et al., PBX1 acts as terminal selector for olfactory bulb dopaminergic neurons. Development 147, (2020).

56. N. Hamada et al., Role of the cytoplasmic isoform of RBFOX1/A2BP1 in establishing the architecture of the developing cerebral cortex. Mol Autism 6, 56 (2015).

57. H. Y. Wang et al., RBFOX3/NeuN is Required for Hippocampal Circuit Balance and Function. Sci Rep 5, 17383 (2015).

58. M. Jacko et al., Rbfox Splicing Factors Promote Neuronal Maturation and Axon Initial Segment Assembly. Neuron 97, 853–868 e856 (2018).

59. J. A. Lee et al., Cytoplasmic Rbfox1 Regulates the Expression of Synaptic and Autism-Related Genes. Neuron 89, 113–128 (2016).

60. X. Dou et al., RBFOX2 recognizes N(6)-methyladenosine to suppress transcription and block myeloid leukaemia differentiation. Nat Cell Biol 25, 1359–1368 (2023).

61. A. Dobin et al., STAR: ultrafast universal RNA-seq aligner. Bioinformatics 29, 15–21 (2013).

62. S. Shen et al., rMATS: robust and flexible detection of differential alternative splicing from replicate RNA-Seq data. Proc Natl Acad Sci U S A 111, E5593–5601 (2014).

63. J. W. Park, S. Jung, E. C. Rouchka, Y. T. Tseng, Y. Xing, rMAPS: RNA map analysis and plotting server for alternative exon regulation. Nucleic Acids Res 44, W333–338 (2016).

64. D. W. Huang et al., DAVID Bioinformatics Resources: expanded annotation database and novel algorithms to better extract biology from large gene lists. Nucleic Acids Res 35, W169–175 (2007).

65. R. Patro, G. Duggal, M. I. Love, R. A. Irizarry, C. Kingsford, Salmon provides fast and bias-aware quantification of transcript expression. Nat Methods 14, 417–419 (2017).

66. M. I. Love, W. Huber, S. Anders, Moderated estimation of fold change and dispersion for RNA-seq data with DESeq2. Genome Biol 15, 550 (2014).

67. A. Shah, Y. Qian, S. M. Weyn-Vanhentenryck, C. Zhang, CLIP Tool Kit (CTK): a flexible and robust pipeline to analyze CLIP sequencing data. Bioinformatics 33, 566–567 (2017).

68. T. Saito, In vivo electroporation in the embryonic mouse central nervous system. Nat Protoc 1, 1552–1558 (2006).

69. S. Assimacopoulos, T. Kao, N. P. Issa, E. A. Grove, Fibroblast growth factor 8 organizes the neocortical area map and regulates sensory map topography. Journal of Neuroscience 32, 7191–7201 (2012).

70. J. C. F. de Winter, Using the Student’s t-test with extremely small sample sizes. *Practical Assessment*, Research, and Evaluation 18, 10 (2013).

71. S. M. Weyn-Vanhentenryck et al., HITS-CLIP and integrative modeling define the Rbfox splicing-regulatory network linked to brain development and autism. Cell Rep 6, 1139–1152 (2014).

