## Supplemental file for "Cell-type-specific splicing of transcription regulators and *Ptbp1* by *Rbfox1/2/3* in the developing neocortex"

#### Figure S1

a

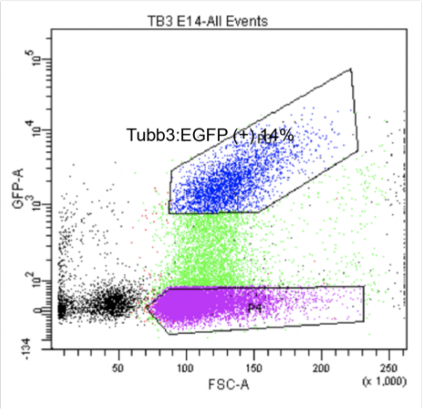

c

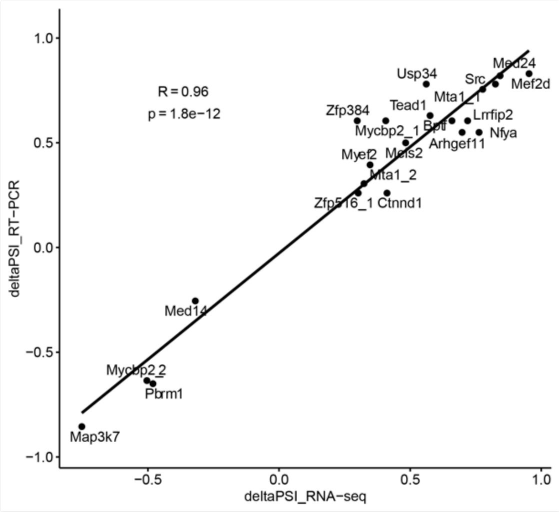

b

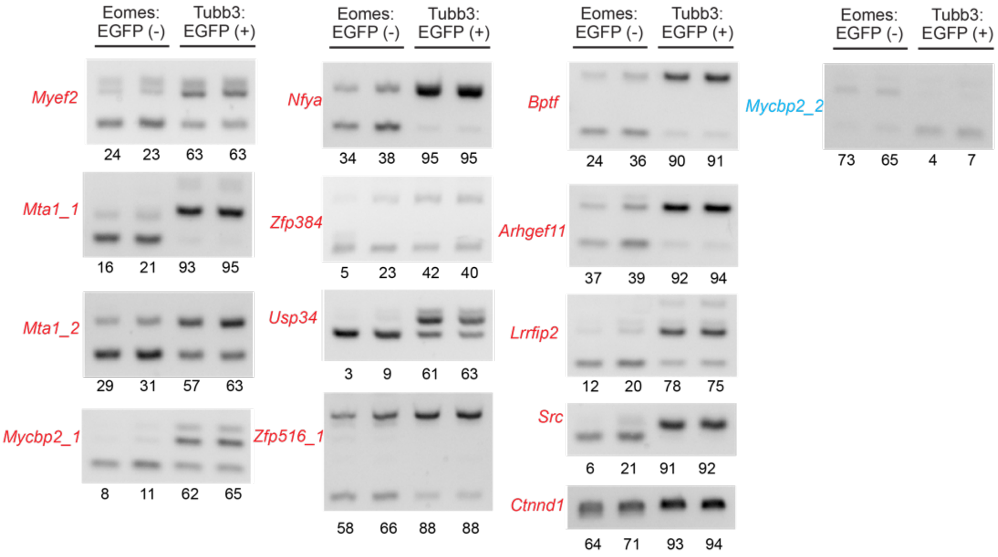

d

| Cell type | Reads | Number of cells |
|---|---|---|
| Neurons | 248299 | 146 |
| Immature.neuron | 137917 | 126 |
| Interneuron | 26968 | 33 |
| IPC | 85084 | 71 |
| RGC | 111232 | 81 |
| Layer.I | 15234 | 10 |
| OPC | 12646 | 9 |
| other | 6510 | 4 |
| Sum | 643890 | 626 |

e

| | | |
|---|---|---|
| Raw data | 3,266,846 | 100.00 |
| Remove 5' and 3' Primers | 1,880,262 | 57.56 |
| Detect UMIs and Cell Barcodes | 1,880,112 | 99.99 |
| Remove PolyA Tail and Artificial Concatemers | 1,758,590 | 93.54 |
| Cluster Reads by Unique Founder Molecules | 1,387,763 | 78.91 |
| Assign to Annotated Cell Types | 643,890 | 46.40 |

f

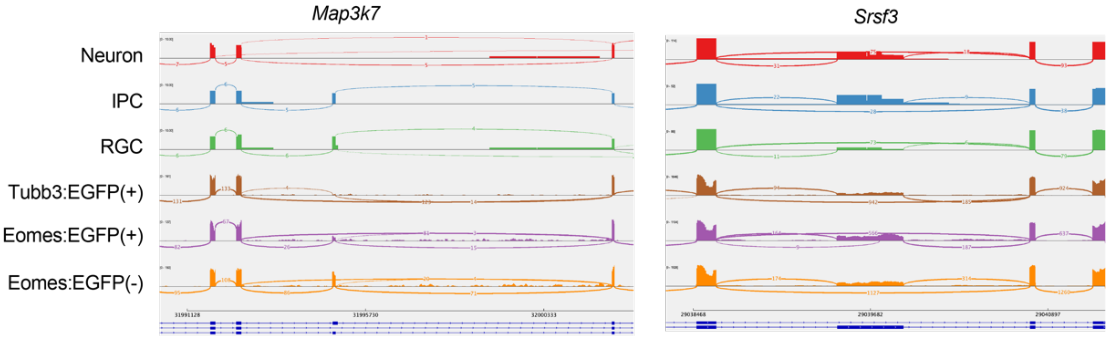

**Figure S1. Transcription regulators are differentially spliced in the developing neocortex.**

a. FACS isolation of *Tubb3:EGFP*(+) neurons from E14.5 mouse dorsal forebrain.
b. RT-PCR validation of neuron-enriched (red) and RGC-enriched (blue) exons in transcription regulators.
c. A scatter plot showing the correlation of ΔPSI values between RNA-Seq and RT-PCR results (21 events shown in Fig. 1f and S1b).
d. Numbers of cells and long reads for each annotated cell type.
e. Numbers and proportions of PacBio Sequel2 reads after each step of filtering and processing.
f. Sashimi plots showing alternative spliced exons in *Map3k7* (included in RGCs and IPCs) and *Srsf3* (enriched in IPCs) in scIso-Seq and FACS-sorted bulk samples.

### Figure S2

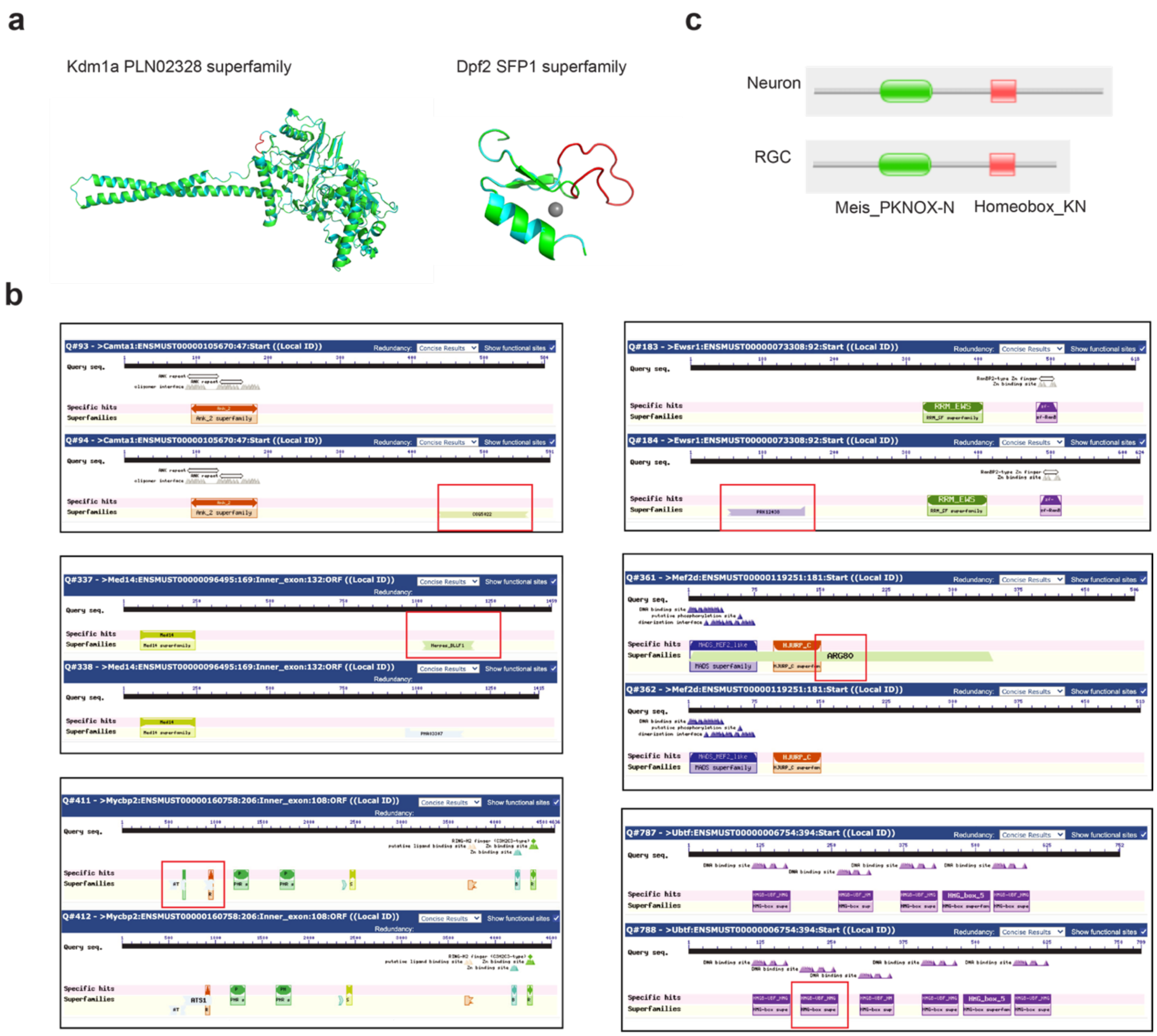

**Figure S2 Differential splicing of transcription regulators affects predicted protein domains.**

a. 3-D protein structure (SWISS-MODEL) showing that alternative splicing of a 12-bp alternative exon in Kdm1a, and a 42-bp SE in Dpf2 inserts extra amino acids into corresponding protein domains. The Gray ball represents a Zinc ion.
b. Paired-wise comparison of protein domains between isoforms (Entrez Conserved Domains Database). The Camta1 21-bp SE following 474 AA is part of a COG5022 superfamily domain; The Ewsr1 18-bp SE following 75 AA is a portion of the PRK12438 superfamily domain; Exclusion of a 132-bp SE at 1049 AA position in Med14 changed the Herpes_BLLF1 superfamily domain; The 21bp Mef2d SE at 285 AA is a part of the ARG80 superfamily domain; Skipping of a108-bp SE in Mycbp2 ENSMUST00000160758 disrupted an ATS1 superfamily domain; The 111-bp SE following 200 AA in Ubtf (ENSMUST00000006754) creates an extra HMGB-UBF_HMG-box.
c. Alternative splicing of *Meis2* is predicted to introduce a longer protein tail in neurons.

#### Figure S3

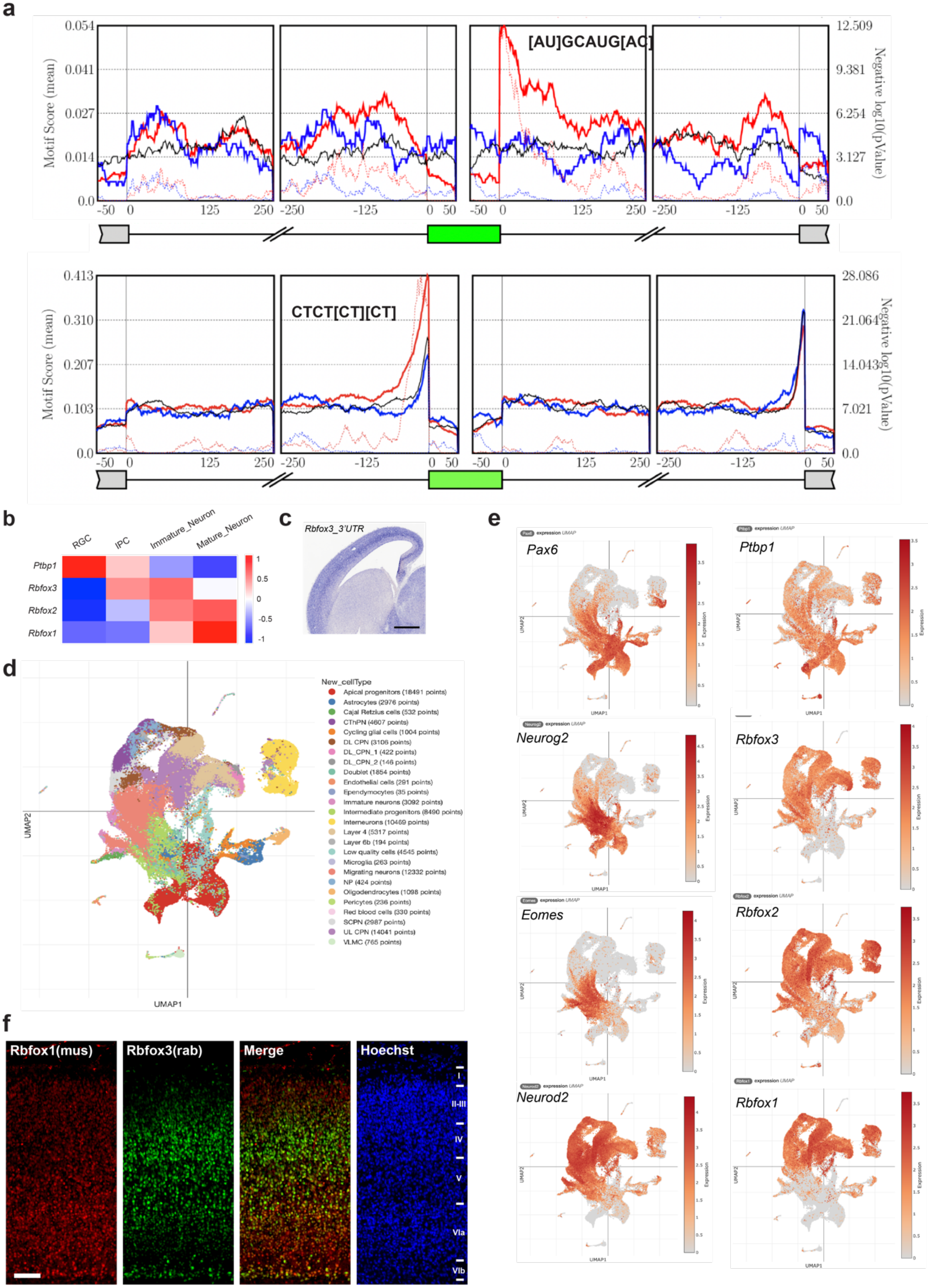

**Figure S3. Differential Rbfox3/2/1 expression in the developing neocortex.**

a. rMAPS analysis of neuron-specific exons showing that the *GCAUG* (Rbfox) motif was significantly enriched in the downstream intron and that the *CTCT[CT][CT]* (Ptbp) motif was significantly enriched in the upstream intron. The green box indicates the alternative exon and position 0 indicates the intron-exon (or exon-intron) boundary. Red lines indicate enrichment scores, and the dashed red lines indicate the p-value (negative log10).
b. Re-analysis of an independent scRNA-Seq dataset(*47*) showing transcription levels of *Ptbp1* and *Rbfox3/2/1* in E14.5 RGCs, IPCs, immature and mature neurons.
c. RNA ISH results of E14.5 sagittal brains showing *Rbfox3* expression with a probe against its 3'UTR. Scale bar, 400μm.
d. Annotated cell types in an independent scRNA-Seq dataset(*47*) that include cells spanning E10.5-P4.
e. Feature plots showing expression levels of *Ptbp1*, *Rbfox3/2/1*, along with *Pax6* (RGC marker), *Eomes* (IPC marker), *Neurog2*, and *Neurod2* (projection neurons).
f. Co-immunostaining results of Rbfox1 (Millipore MABE159, mouse) and Rbfox3 (Cell Signaling Technology 24307T, rabbit) in the P0 mouse neocortex. Scale bar, 100μm.

#### Figure S4

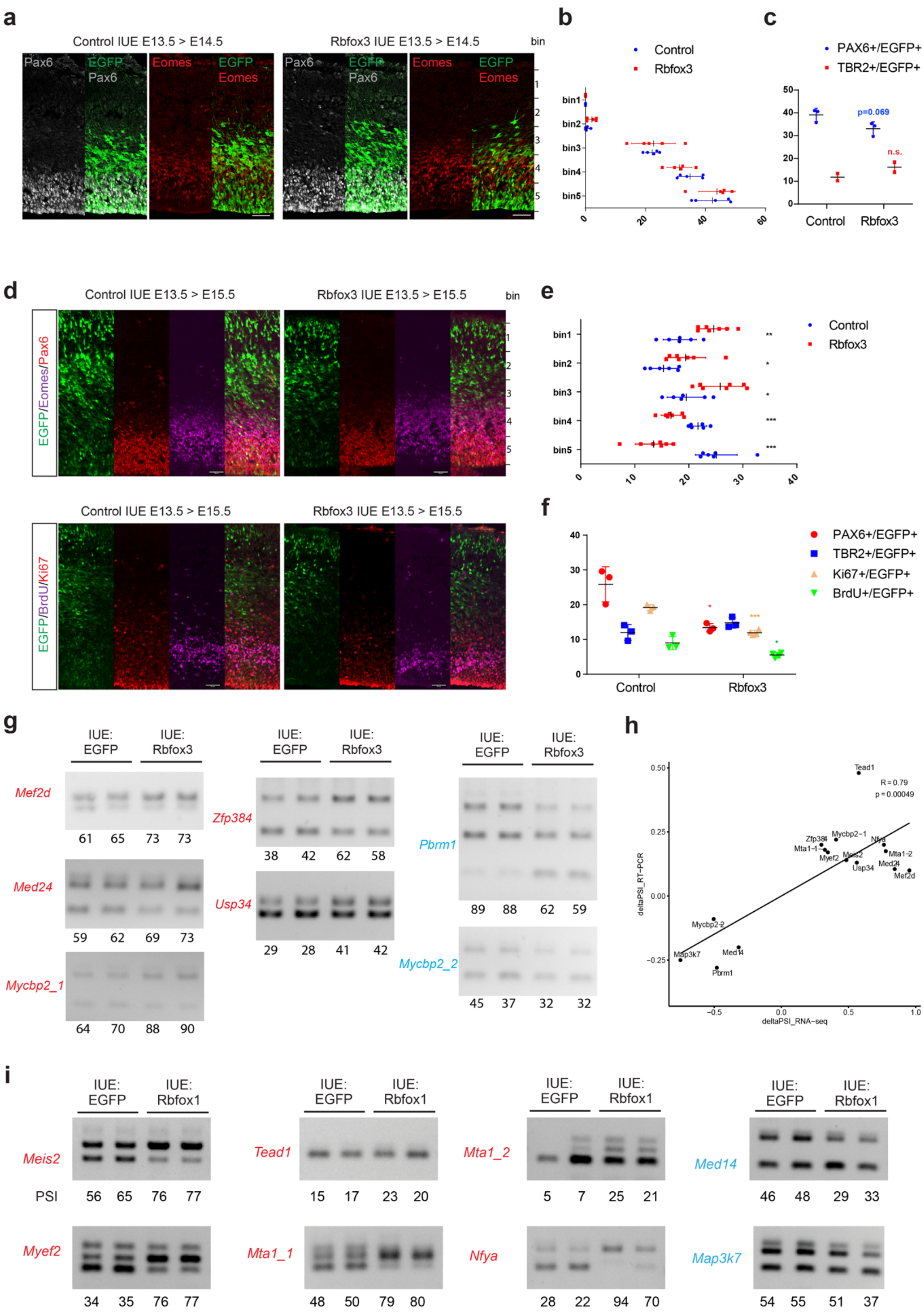
a
Control IUE E13.5 > E14.5
Rbfox3 IUE E13.5 > E14.5
bin
Pax6
EGFP
Eomes
b
Control
Rbfox3
bin1
bin2
bin3
bin4
bin5
c
PAX6+/EGFP+
TBR2+/EGFP+
p=0.069
n.s.
Control
Rbfox3
d
Control IUE E13.5 > E15.5
Rbfox3 IUE E13.5 > E15.5
EGFP/Eomes/Pax6
EGFP/BrdU/Ki67
e
f
PAX6+/EGFP+
TBR2+/EGFP+
Ki67+/EGFP+
BrdU+/EGFP+
g
IUE: EGFP
IUE: Rbfox3
Mef2d
61 65 73 73
Med24
59 62 69 73
Mycbp2_1
64 70 88 90
Zfp384
38 42 62 58
Usp34
29 28 41 42
Pbrm1
89 88 62 59
Mycbp2_2
45 37 32 32
h
R = 0.79
p = 0.00049
deltaPSI_RT−PCR
deltaPSI_RNA−seq
i
IUE: EGFP
IUE: Rbfox1
Meis2
PSI
56 65 76 77
Tead1
15 17 23 20
Mta1_2
5 7 25 21
Med14
46 48 29 33
Myef2
34 35 76 77
Mta1_1
48 50 79 80
Nfya
28 22 94 70
Map3k7
54 55 51 37

**Figure S4. Rbfox proteins switch transcription regulators to neuronal isoforms.**

a. Transient 24-hour expression of empty vector or Rbfox3 in the RGCs, IUE at E13.5 and examination of brain tissues at E14.5. Antibodies against EGFP (green), Pax6 (grey), and Eomes (red). Scale bar, 50 μm.
b. Quantification of the distribution of EGFP-positive cells in the embryonic neocortex (Fig. S4a, unpaired two-tailed Student's t-test).
c. Quantification of Pax6- and Eomes-positive cells in EGFP-positive cells in control and Rbfox3 IUE brain slices (Fig. S4a, unpaired two-tailed Student's t-test).
d. Transient 48-hour expression of empty vector or Rbfox3, IUE at E13.5 and examination of brain tissues at E15.5. BrdU (50 mg/kg) was intraperitoneally injected 30 min before dissection. Scale bar, 50 μm.
e. Quantification of the distribution of EGFP-positive cells in the embryonic neocortex (unpaired two-tailed Student's t-test).
f. Proportions of Pax6-, Eomes-, Ki67- and BrdU-positive cells in EGFP-positive cells in control and Rbfox3 brain slices (unpaired two-tailed Student's t-test).
g. RT-PCR validation of Rbfox3-mediated regulation of neuron- and RGC-specific exons in transcription regulators, numbers indicate PSI values.
h. Correlation of ΔPSI values between RT-PCR (Fig. 4g and S4a) and RNA-Seq data of 15 AS events regulated by Rbfox3.
i. RT-PCR validation of Rbfox1-mediated regulation of neuron- and RGC-specific exons in transcription regulators, numbers indicate PSI values.

### Figure S5

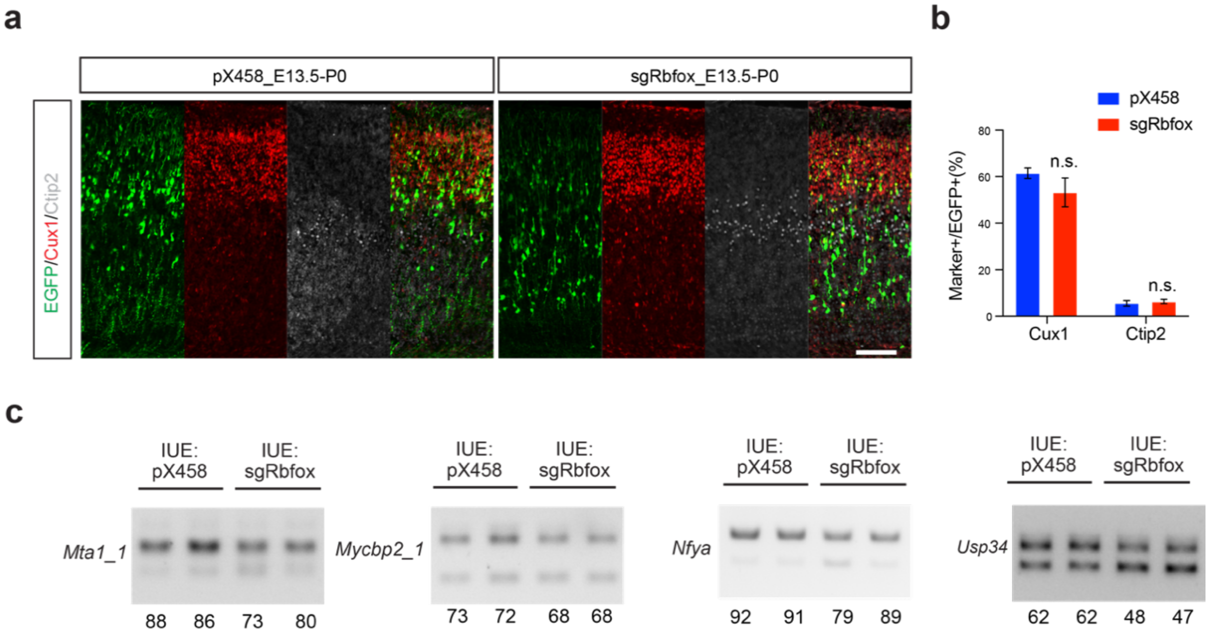

**Figure S5. Rbfox proteins are required for transcription regulators to express neuronal isoforms.**

a. Rbfox1/2/3 triple deletion (green cells) in the neocortex leads to neuronal migration defects; scale bar, 100μm.
b. Quantification of Rbfox1/2/3 triple knockout cells showing that the Cux1- and Ctip2-positive portions were not significantly changed (unpaired two-tailed Student's t-test, n = 4 for each group).
c. RT-PCR results showing that *Rbfox1/2/3* triple knockout decreased the inclusion of neuronal exons in transcription regulators. IUE at E13.5 and analyzed at P0.

### Figure S6

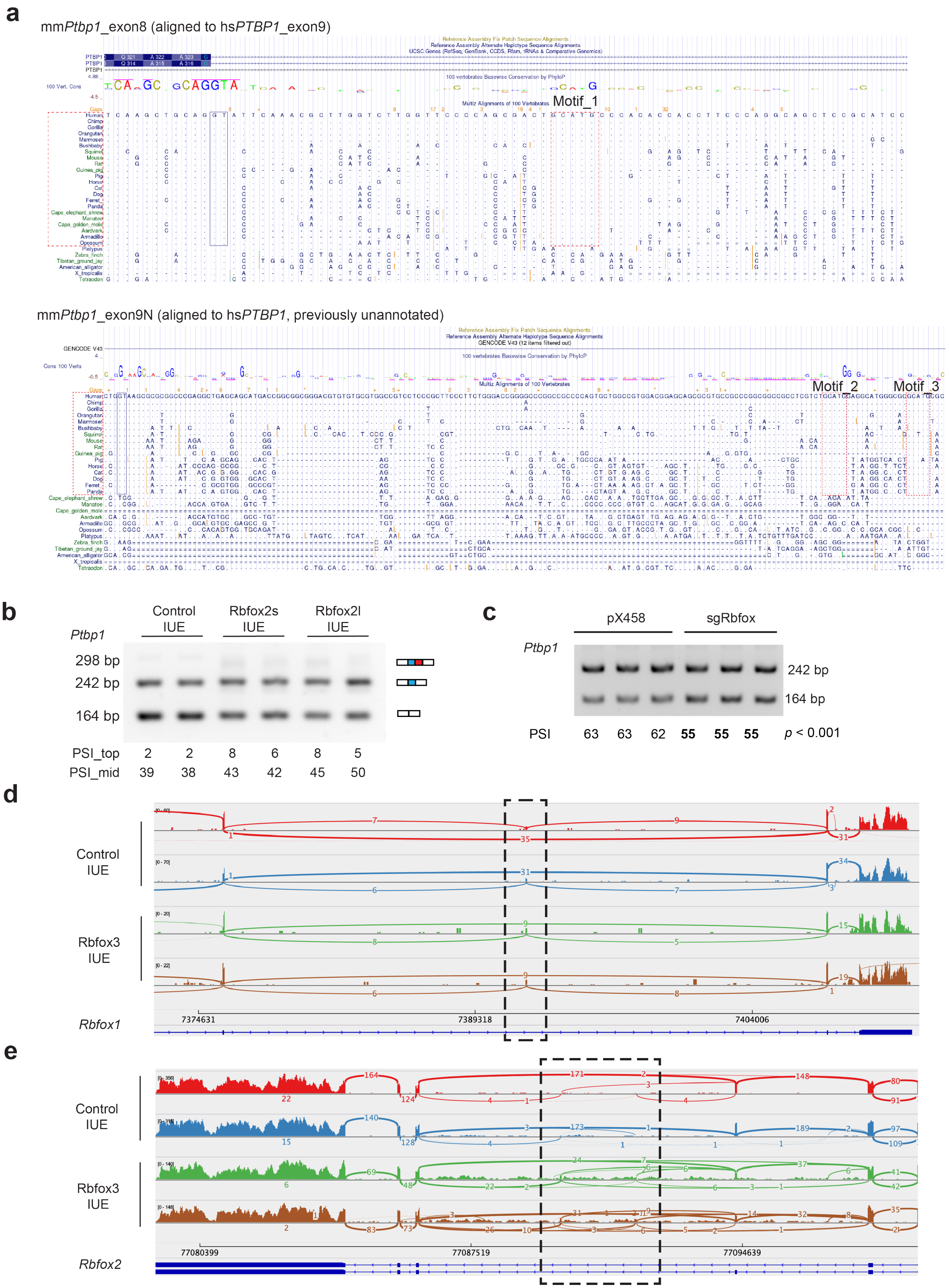

**Figure S6. Crosstalk between *Rbfox* genes and *Ptbp1*.**

a. Multiple sequence alignments across vertebrates showing three conserved *GCAUG* Rbfox binding motifs in mammals.
b. RT-PCR results showing that transient Rbfox2 expression in RGCs promoted *Ptbp1* exon8 and exon9N inclusion.
c. RT-PCR result showing *Ptbp1* exon8 inclusion was decreased in Rbfox1/2/3 triple knockout in Neuro2a cells.
d. Rbfox3 expression induced frame-shift exon inclusion in *Rbfox1*.
e. Rbfox3 transient expression induced noisy splicing in *Rbfox2*.

#### Figure S7

a

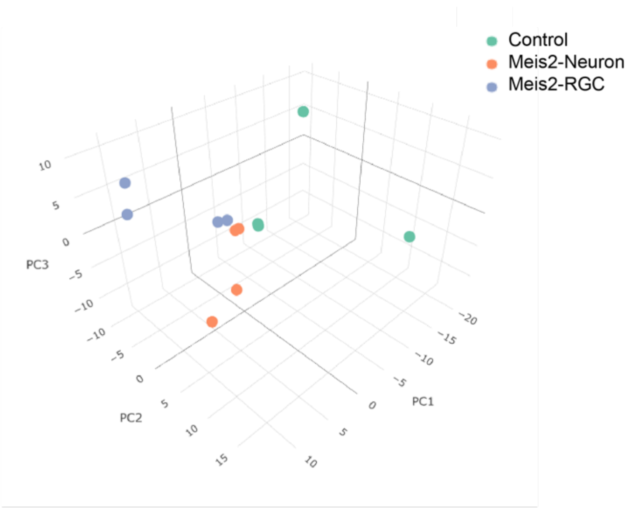

b

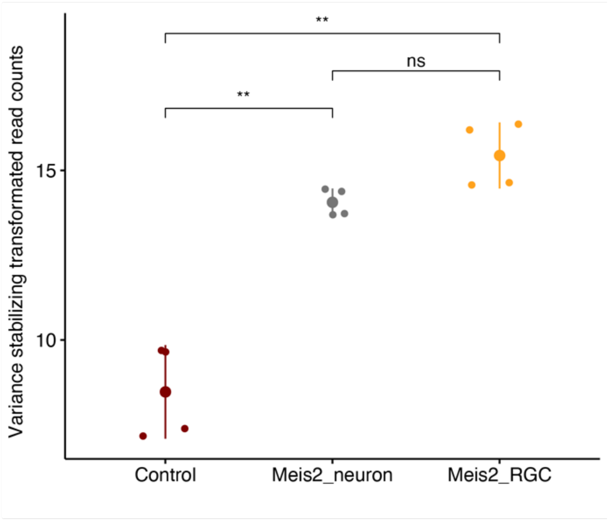

c

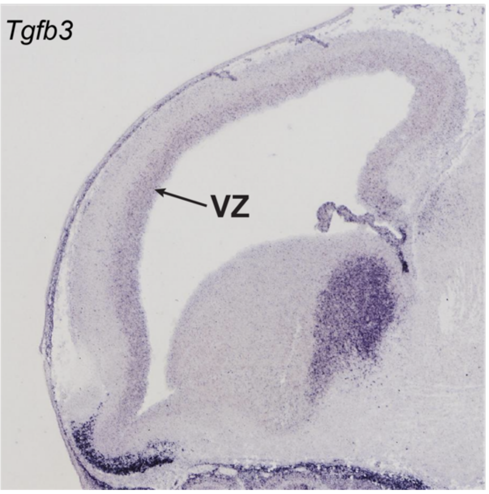

**Figure S7. Meis2 isoforms differentially regulate *Tgfb3* expression.**

a. Principle component analysis of control and *Meis2* RNA-Seq samples.
b. RNA-Seq results showing that ectopic expression of *Meis2* isoforms at comparable levels (statistically not significant, ns). Differential gene expression was determined with the Wald test, corrected by the Benjamini and Hochberg method.
c. RNA ISH results showing that *Tgfb3* was enriched in the ventricular zone (VZ) at E14.5. The images were adapted from Genepaint(*48*).

#### Table S1. Primers for cloning and AS validation

| Name | Sequence |
|---|---|
| CH438 | GGGGGCGGAATTTACGTAGC |
| CH687 | TGGCCTTTTGCTCACATGTTTAACCCTAGAAAGATAGTCTGCGTAAAATTGACGCATGCATTCTTGAAATATTGCTCTCTCTTTCTAAATAGCGCGAATCCGTCGCTGTGCATT<br>TAGGACATCTCAGTCGCCGCTTGGAGCTCCCGTGAGGCGTGCTTGTCAATGCGGTAAGTGTCACTGATTTTGAACTATAACGACCGCGTGAGTCAAAATGACGCATGATTAT<br>CTTTTACGTGACTTTTAAGATTTAACTCATACGATAATTATATTGTTATTTCATGTTCTACTTACGTGATAACTTATTATATATATATTTTCTTGTTATAGATATCACCACTGTAAGA<br>CGGAGAATCTGAACTAGTGAGGGCCTATTTCCCATGATTCCTTCATATTTGCATATACGATACAAGGCTGTTAGAGAGATAATTGGAATTAATTTGACTGTAAACACAAAGATA<br>TTAGTACAAAATACGTGACGTAGAAAGTAATAATTTCTTGGGTAGTTTGCAGTTTTAAAATTATGTTTTAAAATGGACTATCATATGCTTACCGTAACTTGAAAGTATTTCGATTT<br>CTTGGCTTTATATATCTTGTGGAAAGGACGAAACACCTGGCCGGTGTACTCTGGCGCGTTTTAGAGCTAGAAATAGCAAGTTAAAATAAGGCTAGTCCGTTATCAACTTGAAA<br>AAGTGGCACCGAGTCGGTGCTTTTTTGTTTTGAATCGTAAACGTGATCAATGGAACTCGAGGA |
| CH690 | GAATCGTAAACGTGATCAATGGAACTCGAGGAGGGCCTATTTCCCATGAT |
| CH691 | CGCGCCGCCAATGTTAACGCTGTGACCACTCTAAAACAAAAAAGCACCGACTCGGT |
| CH692 | AGTGGTCACAGCGTTAACATTGGCGGCGCGCCGAGGGCCTATTTCCCATGAT |
| CH693 | CGGGCCATTTACCGTAAGTTATGTAACGGGTACCCTAAAACAAAAAAGCACCGACTCGGT |
| CH986 | TCTCATCATTTTGGCAAA GAATTCGCCACCATGTACCCATACGACGTCCCAGACTACGCTGGCGCGCCA TCT AAGTTTGGCACCGTCCT |
| CH987 | GGGCGGAATTTACGTA GCGGCCGC TTA AAGGACAGAATTCCCAGCA |
| CH988 | CAGCCAGCCTTCACTAGACC |
| CH989 | AAAGGACAGAATTCCCAGCA |
| CH990 | CAGTTATGGGGCTGTGTGGGAGGCGCTGGGGTCAGGATGA |
| CH991 | TCATCCTGACCCCAGCGCCTCCCACACAGCCCCATAACTG |
| CH992 | GATCTGCCCGGCTGTGCACG TGCTCCGGCGGCACGCGGAG |
| CH993 | CTCCGCGTGCCGCCGGAGCA CGTGCACAGCCGGGCAGATC |
| XR244 | AGCCAAGGAGCAGCGTATAG |
| XR245 | GGCTTGGCAAATATGAATGC |
| XR293 | ACGATGACGACAAGGGCGCGCCTGCGCAAAGGTACGATGAGCTG |
| XR294 | GGGCGGAATTTACGTAGCGGCCGCTTACATATAGTGCCACTGCCCATCCAT |
| XR296 | GGGCGGAATTTACGTAGCGGCCGCCTATTGGGCATGAATGTCCATAACCTGT |
| XR301 | ACGATGACGACAAGGGCGCGCCTAGCCAAGGAGCAGCGTATAGTCCAG |
| XR302 | GGGCGGAATTTACGTAGCGGCCGCGTGGGGATGGCTTGGCAAATATGAATGCA |
| XR303 | GTATGAATATGGGTATGGATGGGCAGTGGCACTATATGTAACCTTC |
| XR304 | ATCCATACCCATATTCATACCCATTCCACTCATGGGTCCTAGAAGG |
| XR902 | ACACCAATTACTGCTTCCGC |
| XR903 | CTCTGGGTTCAGGGTGTTGT |
| XR1038 | acatttaggtgacactatagaagCTCCGCCTCAGAATGGAAT |
| XR1039 | agtaatacgactcactatagggTGGTGCAGCTCGAAATGTAT |
| XR1040 | acatttaggtgacactatagaaGCTTCCGGCTACAAAGAACA |
| XR1041 | agtaatacgactcactatagGGCTCCGTACAGAGGACAGA |

#### Table S2. Primers for AS validation

| Gene Symbol | target coordinate | forward primer | reverse primer | SE_ Length(bp) | PCR_skipping_ product size(bp) | PCR_inclusion product size(bp) |
|---|---|---|---|---|---|---|
| Mef2d | +chr3:88161780-88161801 | GTCATCCCTGCCAAGTCTCC | CGAGTGGGTAGACTGGGAGA | 21 | 162 | 183 |
| Med24 | -chr11:98717708-98717765 | CTTGCAGTGCCTGGAGAAGA | TGCTGAGATTGGCCAGGATC | 57 | 140 | 197 |
| Src | +chr2:157458842-157458860 | TCACGGACAGAGACTGACCT | AGTTGCTGGGGATGTAACCG | 18 | 127 | 145 |
| Mta1_1 | +chr12:113133540-113133576 | CAGTGTGCTCAGCAGTCTGA | ATGTGCCTGGTCTGTCCATG | 36 | 138 | 174 |
| Nfya | -chr17:48398981-48399068 | AGCTGCCTGGGATCTGTAGA | TGTACCATGATGGGTTGGCC | 87 | 205 | 292 |
| Lrrfip2 | +chr9:111213812-111213914 | AGAAACACATGTGCAGCGTG | CAGCTTCCTGAGACACAGCA | 102 | 177 | 279 |
| Arhgef11 | +chr3:87695397-87695517 | CCTGAAGTCCAAAAGCACGC | CTTCACATGTTCTCTGGCTGG | 120 | 115 | 235 |
| Bptf | -chr11:107086711-107086897 | GGCAGAGAAGAAGGTGGCAT | TGAGTTTGCTCTCCGGGTTC | 186 | 161 | 347 |
| Tead1 | +chr7:112856052-112856064 | TCAGGTTCTTGCCAGAAGGA | AGCTTGTTGTGGATGGCAGT | 12 | 138 | 150 |
| Mycbp2_1 | -chr14:103177255-103177330 | CGGTTACACTGAAGCCTGGT | CATTGTTGCTTCCTGTGGGC | 75 | 139 | 214 |
| Usp34 | +chr11:23362576-23362612 | ACACTGTACTTGGCATCCATGT | TCAGTGTCAGACAGTCGCTG | 36 | 260 | 296 |
| Meis2 | -chr2:115866809-115866904 | TCAGCAACACATGGGGATCC | TCCATAACCTGTCCGCCAAC | 95 | 300 | 395 |
| Ctnnd1 | -chr2:84612531-84612549 | GTTTGCCTCCTCCGGAACTT | TGCTGGATCCTCTGTAGGCT | 18 | 165 | 183 |
| Zfp516_1 | +chr18:82994469-82994537 | AAGGACTTCGCCACCCTCTA | GTGAGGCCACAGTCAACACT | 68 | 171 | 359 |
| Myef2 | -chr2:125098796-125098847 | TGGAGGCATTGGAATGGGAC | ACCAAAGCCTGCCATGCTAT | 51 | 145 | 196 |
| Mta1_2 | +chr12:113120210-113120261 | AGAAGGCGGGACATTTCCAG | TGCCGAGACAGGAACAGTTC | 51 | 149 | 200 |
| Zfp384 | +chr6:125031670-125031853 | CGGAGATGCAGATCCACTCC | TGAAAGCCTTCTCACAGCCC | 183 | 249 | 432 |
| Pbrm1 | +chr14:31113848-31114004 | GACTGGTGGGGACAGAATGG | CCATACGGACTTCCACCTGG | 165 | 121;277 | 286;442 |
| Med14 | -chrX:12689328-12689460 | ATTTCCAAAGCAGCCAGGGA | CAGTTTCCTGCTCGTCCACT | 132 | 171 | 303 |
| Mycbp2_2 | -chr14:103269357-103269465 | CCCTGTACCGGGGGTTAATA | GGTGGTGATTTTGCTTGCGT | 108 | 145 | 253 |
| Map3k7 | +chr4:31994873-31994954 | GAATCTGGACGCCTGAGCTT | CTGACCAGGTTCTGTCCCAG | 81 | 192 | 273 |
| Rnf14 | +chr18:38301655-38301807 | AATGTCGGCAGAAGACCTGG | CACTCTGCCTCGGTGTTCTT | 152 | 212 | 364 |
| Ccar2 | -chr14:70142924-70143088 | GAGCATCCTCTGAAGCAGCT | CCCTGATGCCTCATGTGGTT | 164 | 196 | 360 |
| Mycbp2 | -chr14:103252408-103252526 | ACCTTTGGATATGGGCAGCA | GAAAGTCTGGTCCCCACTCG | 118 | 182 | 300 |
| Crtc2 | +chr3:90262913-90262980 | TGCCTACTGACCAGCGATTG | TTCTGGTGGTTGAGGACGTG | 67 | 376 | 443 |
| Taf1a | +chr1:183405895-183405962 | TCAAGATGGAGCCCGAGAGA | CATCCAGCGAAGTCCAGGAC | 67 | 203 | 270 |
| Ylpm1 | +chr12:85060262-85060314 | TGCAGAAGAAGAGGAAAGCGA | TCGTCAGGGAGCTGAAGGTA | 52 | 119 | 171 |
| Mycbp2_2 | -chr14:103269357-103269465 | CCCTGTACCGGGGGTTAATA | GGTGGTGATTTTGCTTGCGT | 108 | 145 | 253 |
| Map3k7 | +chr4:31994873-31994954 | GAATCTGGACGCCTGAGCTT | CTGACCAGGTTCTGTCCCAG | 81 | 192 | 273 |
| Whsc1 | +chr5:33871406-33871634 | TGAAGAAGCGAAATCGGGCT | TCATCTCCTGGGCTGTCTGA | 228 | 115 | 343 |

#### Table S3. RT-PCR analyses of differential splicing

| **Fig. 1g** | **Emoes:EGFP(-)_rep1** | **Emoes:EGFP(-)_rep2** | **Tubb3:EGFP(+)_rep1** | **Tubb3:EGFP(+)_rep2** | **P_value** | | |
|---|---|---|---|---|---|---|---|
| Meis2 | 31% | 36% | 83% | 85% | 0.0024 | | |
| Tead1 | 7% | 5% | 65% | 72% | 0.0026 | | |
| Map3k7 | 88% | 87% | 4% | 0% | 0.0005 | | |
| Med14 | 75% | 70% | 48% | 46% | 0.0106 | | |
| Mef2d | 10% | 16% | 96% | 96% | 0.0012 | | |
| Med24 | 2% | 5% | 85% | 86% | 0.0006 | | |
| Pbrm1 | 99% | 98% | 27% | 40% | 0.0099 | | |
| **Fig. S1b** | **Emoes:EGFP(-)_rep1** | **Emoes:EGFP(-)_rep2** | **Tubb3:EGFP(+)_rep1** | **Tubb3:EGFP(+)_rep2** | **P_value** | | |
| Myef2 | 24% | 23% | 63% | 63% | 0.0001 | | |
| Mta1_1 | 16% | 21% | 93% | 95% | 0.0009 | | |
| Mta1_2 | 29% | 31% | 57% | 63% | 0.0100 | | |
| Mycbp2_1 | 8% | 11% | 62% | 65% | 0.0018 | | |
| Nfya | 34% | 38% | 96% | 96% | 0.0011 | | |
| Zfp384_rep1 | 5% | 23% | 42% | 40% | 0.0018 | | |
| Zfp384_rep2 | 2% | 22% | 42% | 42% | | | |
| Usp34 | 3% | 9% | 61% | 63% | 0.0026 | | |
| Zfp516_1 | 58% | 66% | 88% | 88% | 0.0239 | | |
| Bptf | 24% | 36% | 90% | 91% | 0.0098 | | |
| Arhgef11 | 37% | 39% | 92% | 94% | 0.0007 | | |
| Lrrfip2 | 12% | 20% | 78% | 75% | 0.0049 | | |
| Src | 6% | 21% | 91% | 92% | 0.0092 | | |
| Ctnnd1 | 64% | 71% | 93% | 94% | 0.0180 | | |
| Mycbp2_2 | 73% | 65% | 4% | 7% | 0.0052 | | |
| **Fig. 2h** | **DMSO_rep1** | **DMSO_rep2** | **DMSO_rep3** | **Cycloheximide_rep1** | **Cycloheximide_rep2** | **Cycloheximide_rep3** | **P_value** |
| Crtc2 | 49% | 43% | 50% | 63% | 68% | 67% | 0.0022 |
| **Fig. 2i** | **DMSO_rep1** | **DMSO_rep2** | **DMSO_rep3** | **Cycloheximide_rep1** | **Cycloheximide_rep2** | **Cycloheximide_rep3** | **P_value** |
| Whsc1 | 13% | 10% | 12% | 21% | 18% | 24% | 0.0086 |
| **Fig. 2j** | **DMSO_rep1** | **DMSO_rep2** | **DMSO_rep3** | **Cycloheximide_rep1** | **Cycloheximide_rep2** | **Cycloheximide_rep3** | **P_value** |
| Ccar2 | 96% | 95% | 95% | 77% | 76% | 74% | 0.0000 |
| Taf1a | 97% | 97% | 96% | 86% | 85% | 86% | 0.0000 |
| Ylpm1 | 97% | 97% | 97% | 81% | 82% | 79% | 0.0001 |
| **Fig. 4g** | **IUE:EGFP_rep1** | **IUE:EGFP_rep2** | **IUE:Rbfox3_rep1** | **IUE:Rbfox3_rep2** | **P_value** | | |
| Meis2 | 59% | 62% | 74% | 75% | 0.0125 | | |
| Myef2 | 47% | 45% | 62% | 64% | 0.0068 | | |
| Tead1 | 10% | 11% | 56% | 61% | 0.0028 | | |
| Mta1_1 | 25% | 32% | 45% | 48% | 0.0420 | | |
| Mta1_2 | 58% | 61% | 75% | 79% | 0.0198 | | |

| | | | | | | | | |
|---|---|---|---|---|---|---|---|---|
| Nfya | 75% | 76% | 94% | 97% | 0.0062 | | | |
| Med14 | 62% | 54% | 36% | 40% | 0.0465 | | | |
| Map3k7 | 67% | 62% | 41% | 38% | 0.0133 | | | |
| **Fig. 4h** | **IUE:EGFP _rep1** | **IUE:EGFP _rep2** | **IUE:Rbfox2s _rep1** | **IUE:Rbfox2s _rep2** | **IUE:Rbfox2l _rep1** | **IUE:Rbfox2l _rep2** | **P_value(Rbfox2s vs EGFP)** | **P_value(Rbfox2l vs EGFP)** |
| Meis2 | 48% | 43% | 51% | 53% | 64% | 59% | 0.1372 | 0.0455 |
| Myef2 | 31% | 35% | 43% | 47% | 55% | 59% | 0.0513 | 0.0136 |
| Tead1 | 22% | 21% | 69% | 60% | 77% | 86% | 0.0109 | 0.0056 |
| Mta1_1 | 48% | 50% | 72% | 72% | 82% | 76% | 0.0019 | 0.0109 |
| Mta1_2 | 31% | 32% | 42% | 37% | 45% | 43% | 0.0883 | 0.0079 |
| Nfya | 65% | 57% | 84% | 87% | 90% | 90% | 0.0291 | 0.0185 |
| Med14 | 43% | 50% | 38% | 31% | 29% | 26% | 0.1362 | 0.0379 |
| Map3k7 | 49% | 57% | 18% | 27% | 19% | 16% | 0.0368 | 0.0142 |
| **Fig. S4g** | **IUE:EGFP _rep1** | **IUE:EGFP _rep2** | **IUE:Rbfox3 _rep1** | **IUE:Rbfox3 _rep2** | **P_value** | | | |
| Mef2d | 62% | 66% | 75% | 76% | 0.0338 | | | |
| Med24_rep1 | 59% | 61% | 69% | 73% | 0.0010 | | | |
| Med24_rep2 | 53% | 61% | 73% | 76% | | | | |
| Mycbp2_1 | 64% | 70% | 88% | 90% | 0.0200 | | | |
| Zfp384 | 38% | 42% | 62% | 58% | 0.0194 | | | |
| Usp34 | 29% | 28% | 41% | 42% | 0.0029 | | | |
| Pbrm1 | 89% | 88% | 62% | 59% | 0.0032 | | | |
| Mycbp2_2_rep1 | 45% | 37% | 32% | 32% | 0.0425 | | | |
| Mycbp2_2_rep2 | 46% | 43% | 40% | 37% | | | | |
| **Fig. 5d** | **pX458 _rep1** | **pX458 _rep2** | **sgRbfox _rep1** | **sgRbfox _rep2** | **P_value** | | | |
| Meis2 | 88% | 87% | 77% | 77% | 0.0019 | | | |
| Myef2_rep1 | 53% | 54% | 38% | 44% | 0.0062 | | | |
| Myef2_rep2 | 52% | 53% | 41% | 49% | | | | |
| Tead1 | 43% | 50% | 17% | 24% | 0.0344 | | | |
| Zfp384_rep1 | 31% | 30% | 19% | 27% | 0.0126 | | | |
| Zfp384_rep2 | 29% | 29% | 15% | 21% | | | | |
| **Fig. S5c** | **pX458 _rep1** | **pX458 _rep2** | **sgRbfox _rep1** | **sgRbfox _rep2** | **P_value** | | | |
| Mta1_1_rep1 | 88% | 86% | 73% | 80% | 0.0105 | | | |
| Mta1_1_rep2 | 92% | 89% | 76% | 85% | | | | |
| Mycbp2_1 | 73% | 72% | 68% | 68% | 0.0167 | | | |
| Usp34 | 62% | 63% | 48% | 47% | 0.0022 | | | |
| Nfya_rep1 | 92% | 91% | 79% | 89% | 0.0423 | | | |
| Nfya_rep2 | 94% | 93% | 80% | 90% | | | | |
| **Fig. 6c** | **Control: IUE _rep1** | **Control: IUE _rep2** | **Rbfox3: IUE _rep1** | **Rbfox3: IUE _rep2** | **P_value** | | | |

| | | | | | | | |
|---|---|---|---|---|---|---|---|
| Ptbp1 | 37% | 39% | 48% | 48% | 0.0096 | | |
| **Fig. 6g** | **PSI_top _Rep1** | **PSI_top _Rep2** | **PSI_mid _Rep1** | **PSI_mid _Rep2** | **P_value _top** | **P_value _mid** | |
| WT | 0% | 0% | 37% | 39% | | | |
| WT+Rbfox3 | 23% | 29% | 43% | 38% | 0.0131 | 0.6707 | |
| Del_Motif_1 | 0% | 0% | 40% | 40% | | | |
| Del_Motif_1 +Rbfox3 | 26% | 26% | 37% | 37% | #DIV/0! | #DIV/0! | |
| Del_Motif _2-3 | 0% | 0% | 45% | 44% | | | |
| Del_Motif_2-3+Rbfox3 | 0% | 0% | 61% | 60% | | 0.0019 | |
| Del_Motif_ 1-2-3 | 0% | 0% | 55% | 53% | | | |
| Del_Motif _1-2-3+Rbfox3 | 0% | 0% | 51% | 52% | | 0.1548 | |
| **Fig. S6b** | **PSI_top _Rep1** | **PSI_top _Rep2** | **PSI_mid _Rep1** | **PSI_mid _Rep2** | **P_value _top** | **P_value _mid** | |
| Control IUE | 2% | 2% | 39% | 38% | | | |
| Rbfox2s IUE | 8% | 6% | 43% | 42% | 0.0377 | 0.7655 | |
| Rbfox2l IUE | 8% | 5% | 45% | 50% | 0.0955 | 0.0539 | |
| **Fig. S6c** | | | | | | | |
| **sgRbfox3 in Neuro2a cells** | **pX458 _rep1** | **pX458 _rep2** | **pX458 _rep3** | **sgRbfox _rep1** | **sgRbfox _rep2** | **sgRbfox _rep3** | **P_value** |
| Ptbp1 | 63% | 63% | 62% | 55% | 55% | 55% | 0.0001 |
| **Fig. 7c** | | | | | | | |
| **Meis2 minigene** | **WT _rep1** | **WT _rep2** | **Ex12_mut _rep1** | **Ex12_mut _rep2** | **Del_1 _rep1** | **Del_1 _rep2** | |
| | 91% | 89% | 46% | 48% | 81% | 81% | |
| **P_value** | | | | 0.00108 | | 0.01212 | |
| | **Ex12_mut+Del _1_rep1** | **Ex12_mut+Del _1_rep2** | **Del_2 _rep1** | **Del_2 _rep2** | **Ex12_mut+ Del_2_rep1** | **Ex12_mut+ Del_2_rep2** | |
| | 27% | 31% | 90% | 91% | 35% | 33% | |
| **P_value** | | 0.01508 | | 0.69849 | | 0.01163 | |
| | **Del_3 _rep1** | **Del_3 _rep2** | **Ex12_mut+ Del_3_rep1** | **Ex12_mut+ Del_3_rep2** | | | |
| | 89% | 88% | 48% | 46% | | | |
| **P_value** | | 0.31175 | | 1.00000 | | | |
